# SDC3 acts as a timekeeper of myogenic differentiation by regulating the insulin/AKT/mTOR axis in muscle stem cell progeny

**DOI:** 10.1101/2020.08.10.244152

**Authors:** Fiona K. Jones, Alexander Phillips, Andrew R. Jones, Addolorata Pisconti

## Abstract

Muscle stem cells (MuSCs) are indispensable for muscle regeneration. A multitude of extracellular stimuli direct MuSC fate decisions from quiescent progenitors to differentiated myocytes. The activity of these signals is modulated by coreceptors such as syndecan-3 (SDC3). We investigated the global landscape of SDC3-mediated regulation of myogenesis using a phosphoproteomics approach which revealed, with the precision level of individual phosphosites, the large-scale extent of SDC3-mediated regulation of signal transduction in MuSCs. We then focused on INSR/AKT/mTOR as a key pathway regulated by SDC3 during myogenesis and mechanistically dissected SDC3-mediated inhibition of insulin signaling in MuSCs. SDC3 interacts with INSR limiting insulin signal transduction via AKT/mTOR. Both knockdown of INSR and inhibition of AKT rescue *Sdc3^-/-^* MuSC differentiation to wild type levels. Since SDC3 is rapidly downregulated at the onset of differentiation, our study suggests that SDC3 acts a timekeeper to restrain proliferating MuSC response to insulin and prevent premature differentiation.

## INTRODUCTION

Adult myogenesis includes both muscle growth and regeneration and is carried out by muscle stem cells (MuSCs) in response to hypertrophic stimuli or injury, respectively. MuSCs are quiescent myogenic progenitors that become activated in response to various stimuli and subsequently proliferate and differentiate inside the musculature, ultimately fusing with damaged muscle fibers (myofiber) to repair them, or with one another to generate new myofibers. During this process, MuSCs also self-regenerate, such that a pool of quiescent MuSCs poised to activate on demand is maintained, more or less, throughout life (Olguin and Pisconti, 2012). The mechanisms that regulate myogenesis and MuSC self-renewal are partly intrinsically encoded (Dumont et al., 2015) and partly the result of MuSC response to environmental cues (Mashinchian et al., 2018).

Syndecans, a type of transmembrane heparan sulfate proteoglycans, are multifaceted regulators of cell signaling, as their dual nature of protein and glycan confers them the ability to bind multiple partners simultaneously, thus coordinating the transduction of multiple signaling pathways (Pisconti et al., 2012). The extent to which syndecans contribute to global signaling regulation in MuSC progeny has been recently revealed by single cell RNA-sequencing studies (De Micheli et al., 2020). However, the mechanistic details underlying such powerful regulatory properties of syndecans remain poorly understood.

The four syndecans are dynamically expressed in developing (Cornelison et al., 2001; Liu et al., 2004; Olguin and Brandan, 2001; Pisconti et al., 2012; Velleman and Song, 2017) and regenerating (Casar et al., 2004; Cornelison et al., 2001; Fuentealba et al., 1999; Pisconti et al., 2016; Pisconti et al., 2012; Pisconti et al., 2010; Ronning et al., 2020; Shin et al., 2013) muscle. Based on knockout studies, syndecan-3 (SDC3) and syndecan-4 (SDC4) appear to fulfil indispensable and distinct functions in myogenesis (Cornelison et al., 2004). SDC3 is especially interesting in MuSC biology as its presence is necessary for maintenance of MuSC quiescence and their subsequent progression through the cell cycle once activated (Pisconti et al., 2010). However, SDC3 loss is not detrimental to muscle regeneration or homeostasis, quite the contrary. *Sdc3^-/-^* mice show improved muscle maintenance and regeneration in models of acute injury, aging and muscular dystrophy (Pisconti et al., 2016; Pisconti et al., 2010). We and others have previously shown that this is partly due to SDC3’s ability to regulate Notch and FGF signaling in a coordinated manner, which prevents MuSC exhaustion and effectively enhances regeneration (Pisconti et al., 2010). However, the observation that SDC3 loss induces global changes in phospho-tyrosine levels in MuSCs (Cornelison et al., 2004) suggests that FGF and Notch may represent only the “tip of the iceberg” and that more signaling pathways owe their regulation to SDC3.

We used an unbiased phosphoproteomics approach to interrogate SDC3-dependent signal transduction at a global level and identified insulin signaling mediated by AKT/mTOR as a key pathway regulated by SDC3 in muscle progenitors. We then used biochemistry and cell biology approaches to dissect the insulin pathway in primary and immortalized muscle progenitors, to identify at which level SDC3 regulates it. Lastly, we show how SDC3 expression acts as a molecular switch for the insulin/AKT/mTOR pathway promoting myoblast differentiation.

## RESULTS

### SDC3 depletion in myoblasts leads to increased tyrosine phosphorylation in a cell autonomous manner

Our previous work on SDC3-mediated regulation of Notch signaling strongly suggests that SDC3 loss is responsible for the cell cycle and self-renewal phenotypes observed *in vitro* and *in vivo* in a muscle stem cell (MuSC)-autonomous manner (Pisconti et al., 2010) To definitively test whether SDC3 functions in a cell-autonomous manner in myogenesis, we stably knocked-down SDC3 in C2C12 myoblasts (Figure 1A) and assayed the phenotype of a stable cell line where SDC3 was heavily depleted (Figure 1B, knockdown efficiency ≥95%) by comparison to a stable cell line transfected with an empty vector (from now on referred to as S3^kd^ and Ctrl, respectively). SDC3 depletion in myoblasts led to a phenotype comparable to that of primary *Sdc3^-/-^* MuSC progeny (Cornelison et al., 2004; Pisconti et al., 2016; Pisconti et al., 2010), including: (i) increased responsiveness to FGF2 stimulation (Figure 1C and Figure S1); (ii) reduced proliferation rate (Figure S1); (iii) enhanced differentiation and fusion (Figure 1D-F); and (iv) myotube hypertrophy (Figure 1G). These findings further support the idea that loss of SDC3 affects myogenesis in a way that is largely MuSC-autonomous.

**Figure 1.**
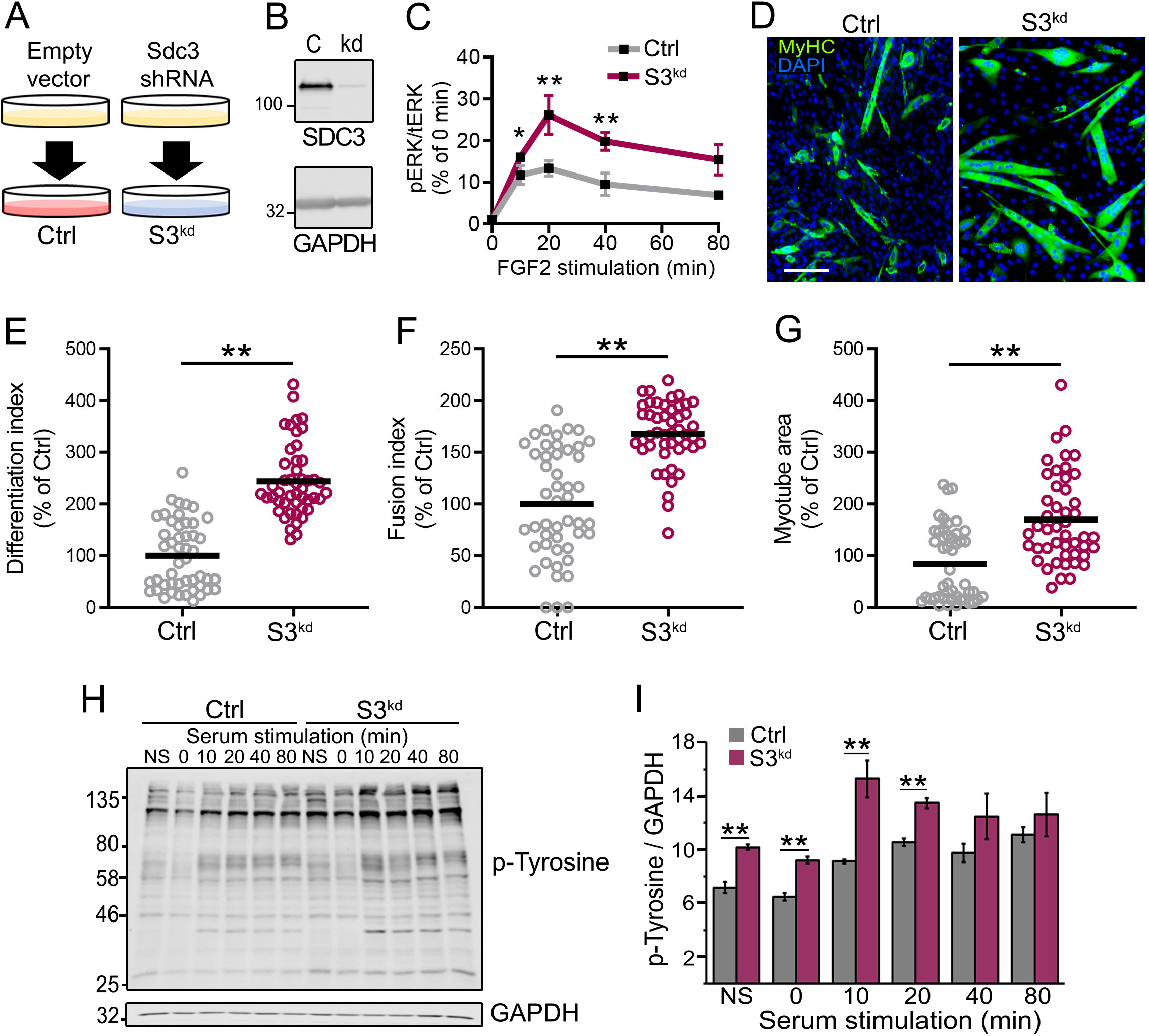
SDC3-depleted cells recapitulate *Sdc3^-/-^* myoblast phenotypes. **(A)** Schematic of the protocol used to stably knockdown SDC3 in C2C12 myoblasts. An empty vector was transfected to generate the control (Ctrl) cell line and Sdc3 shRNA was transfected to generate the SDC3-depleted (S3^kd^) cell line. **(B)** Control **(C)** and S3^kd^ (S3) myoblasts were lysed in RIPA buffer and treated with heparinase III and chondroitinase ABC to remove glycosaminoglycans. Western blotting was used to measure the levels of SDC3 in both cell lines. GAPDH used as loading control. SDC3 is detected as a detergent-resistant dimer by western blotting. (C) Ctrl (gray) and S3^kd^ (purple) myoblasts were serum-starved for five hours then stimulated with 2 nM FGF2 over an 80 minutes time-course. Cells were lysed at the indicated time points and levels of phosphorylated ERK1/2 (pERK) and total ERK1/2 (tERK) quantified via western blotting. All band intensities were quantified using ImageJ. For each of three independent experiments, pERK1/2 band intensities were normalized to total ERK1/2 band intensities at each time point and then each time point expressed as fold change of time 0. In figure, each time point subsequent to time 0 is plotted as average of the 3 experiments (N = 3). **(D)** Immunofluorescent images of Ctrl and S3^kd^ myoblasts induced to differentiate for 5 days by reducing the serum concentration. Cells were fixed then stained to detect myosin heavy chain (MyHC, green, MF20 clone antibody) and DNA (DAPI, blue) to visualize differentiated cells and nuclei respectively. Scale bar = 200 μm. N = 3. **(E-G)** Quantification of differentiation index (E), fusion index (F) and myotube area (G). Ctrl and S3^kd^ myoblasts were cultured, treated and imaged as in (D). Averages from 3 independent experiments are plotted where in each experiment data from S3^kd^ myoblasts are expressed as a percentage of Ctrl myoblasts. ** = p < 0.01. Each biological replicate was plated in triplicate (3 biological replicates x 3 technical replicate per cell line) and 16 images per biological replicate were scored (total N□=□48 images scored per data-point). **(H)** Ctrl and S3^kd^ myoblasts were serum-starved, stimulated with 10% fetal bovine serum and then lysed at various time points over an 80 minutes time course, or lysed prior to serum-starvation (NS). Cells were lysed in RIPA buffer and levels of phosphorylated tyrosines were visualized via western blotting. GAPDH was used as loading control. N = 3, representative western blot images are shown. **(I)** Quantification of (H). Error bars represent S.E.M. ** = p < 0.01. Averages of 3 independent experiments are plotted (N = 3).

One of the most intriguing and yet unexplained phenotypes of primary *Sdc3^-/-^* MuSC progeny is an overall increase in tyrosine phosphorylation (Cornelison et al., 2004), which is suggestive of SDC3 playing a crucial role in the regulation of several signaling pathways simultaneously.

One of the most intriguing and yet unexplained phenotypes of primary *Sdc3^-/-^* MuSC progeny is an overall increase in tyrosine phosphorylation (Cornelison et al., 2004), which is suggestive of SDC3 playing a crucial role in the regulation of several signaling pathways simultaneously. This increased phosphotyrosine phenotype was also recapitulated in S3^kd^ myoblasts, both in serum-starved cells and, especially, in cells re-stimulated with serum (Figure 1H-I). Since serum contains a plethora of signaling molecules (growth factors, cytokines, hormones, etc.), these data further support a role for SDC3 in the coordination of signal transduction via several receptors.

### Phosphoproteomics reveals a role for SDC3 in the regulation of insulin/PI3K/mTOR signaling

To understand which signaling pathways, amongst the several that respond to serum stimulation, are regulated by SDC3 in myoblasts, we undertook a shotgun phosphoproteomics approach. We profiled phosphopeptide abundance in Ctrl and S3^kd^ myoblasts and studied how they change in response to SDC3 depletion and/or serum stimulation, with the aim of identifying candidate signaling pathways to then validate in primary MuSC progeny isolated from wild type and *Sdc3^-/-^* mice. Following phosphopeptide enrichment on titanium dioxide (TiO_2_) columns and label free LC-MS/MS (Figure 2A-C and Jones et al. (2019) we identified and quantified changes in serine, threonine and tyrosine phosphorylation dependent on either SDC3 depletion (comparisons 1 and 2 in Figure 2C), serum stimulation (comparisons 3 and 4 in Figure 2C) or both combined in a two-dimensional analysis (comparison 5 in Figure 2C). We identified a total of 11,777 peptides of which 7,190 were phosphopeptides, corresponding to 1,871 phosphoproteins.

**Figure 2.**
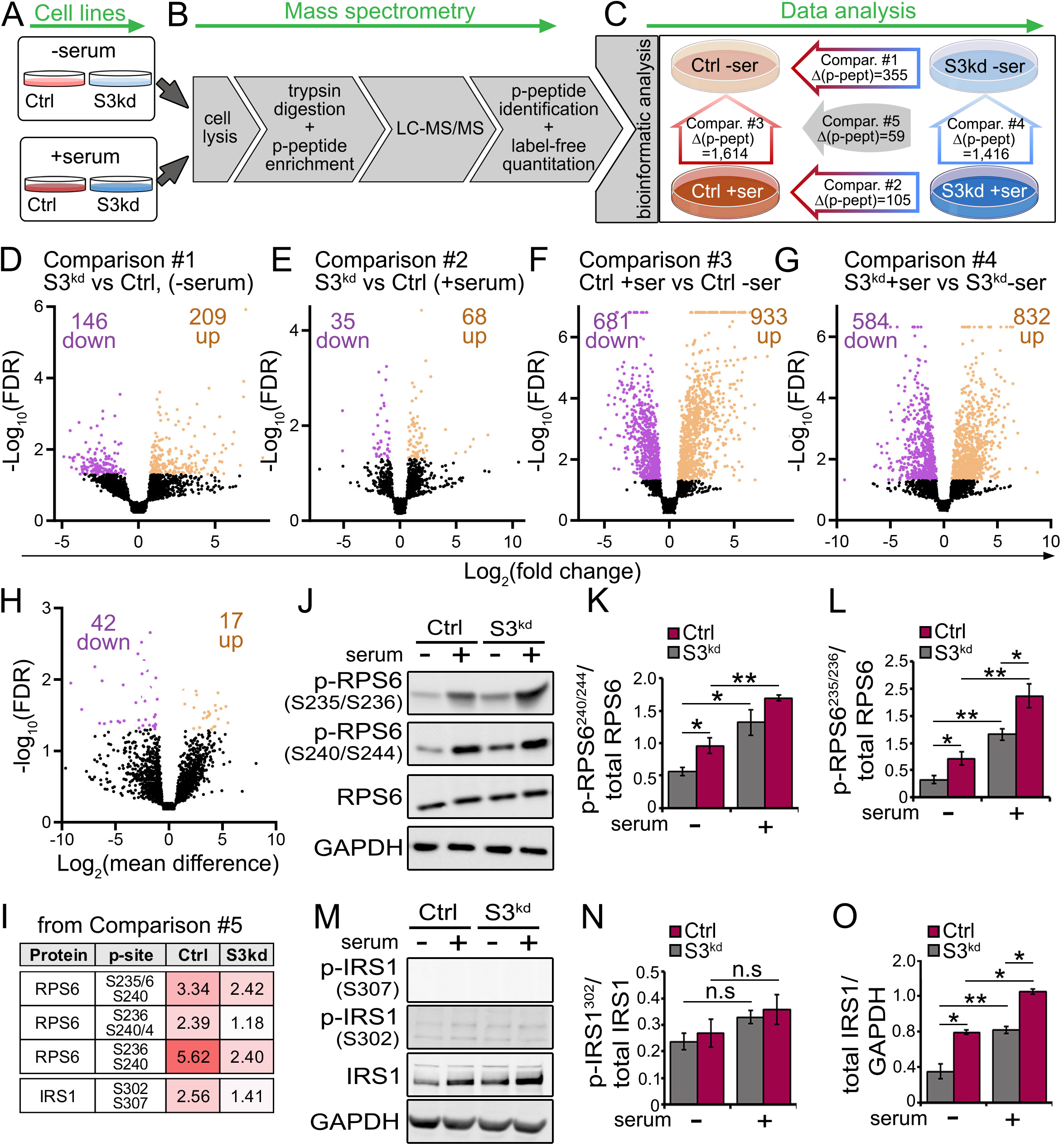
Phosphoproteomics reveals widespread failure of S3^kd^ myoblasts to decrease signal transduction upon serum removal. **(A-C)** Schematic depicting the phosphoproteomics workflow. Control (Ctrl) and SDC3-depleted (S3^kd^) myoblasts were serum-starved for 5 hours before being re-stimulated with 10% fetal bovine serum (FBS) for 10 minutes and then lysed (A). Cell lysates were enriched for phosphopeptides (p-peptide) on TiO_2_ columns and then analyzed via LC-MS/MS. P-peptides were identified and quantified using a label-free method (B). A Bayesian inference model was used to determine statistically significant differences in phosphopeptide abundance across all four datasets, which were compared pairwise in five different ways that generated five lists of differentially abundant peptides: Comparison #1 = differences between S3^kd^ and Ctrl myoblasts at the serum starved state; Comparison #2 = differences between S3^kd^ and Ctrl myoblasts at 10 minutes of serum stimulation; Comparison #3 = differences between serum-starved and serum-stimulated Ctrl myoblasts; Comparison #4 = differences between serum-starved and serum-stimulated S3^kd^ myoblasts; Comparison #5 = p-peptides regulated by both S3^kd^ depletion and serum stimulation. Δ(p-pept) = number of phosphopeptides that are significantly different in abundance between the conditions indicated by the relevant arrow (C). **(D-H)** Volcano plots displaying all phopshopeptides identified in comparisons #1-5. P-peptides were considered significant if q-value < 0.05. N = 4 independent cell cultures per condition. **(I)** Natural log change in phosphopeptide abundance reported from phosphoproteomics data. **(J)** Control (Ctrl) and SDC3-depleted (S3^kd^) myoblasts were serum-starved for 5 hours then re-stimulated with serum for 10 minutes. Cells were lysed and processed for western blotting. Levels of phosphorylated-RPS6^S240/S244^, phosphorylated RPS6^S235/235^ or total RPS6 were measured. GAPDH was measured as a loading control. N = 3. **(K-L)** Quantification of (H) using ImageJ. * = p<0.05, ** = p<0.01. N = 3. **(M)** The same cell lysates used for proteomics analysis shown in (A-H) were used to measure the levels of phosphorylated-IRS1^S307^, IRS1^S302^ and total IRS1 by western blotting. GAPDH was measured as a loading control. N = 3. **(N-O)** Quantification of (K) using ImageJ. * = p<0.05, ** = p<0.01, n.s = non-significant. N = 3.

Firstly, we investigated the effects of SDC3 depletion on the myoblast phosphoproteome in the absence of serum (Figure 2D, Comparison #1) and presence (Figure 2E, Comparison #2) of serum. In both cases, the ratio between upregulated and downregulated phosphopeptides (Up/Down in Figure 2D-E), was >1 and was greater after serum stimulation. This is consistent with our previous results on tyrosine phosphorylation (Figure 1I), where the phospho-tyrosine complement of SDC3-depleted myoblasts was more abundant than that of Ctrl myoblasts, especially after serum stimulation. Intriguingly, SDC3 loss affected the myoblast phosphoproteome ~3 times more in the absence of serum (total Δp-peptides = 355, Figure 2D) than in the presence of serum (Δp-peptides = 105, Figure 2E). The latter finding suggests that SDC3 might control the extent to which a pathway is active so that when the extracellular signal that activates a given pathway is only present at low levels (e.g. the remnants of growth factors bound the extracellular and pericellular matrix after serum washout), the importance of SDC3 is more evident than when the same signal is present at saturating levels.

Secondly, we investigated myoblast response to serum stimulation in the presence (Figure 2F, Comparison #3) and near-absence (Figure 2G, Comparison #4) of SDC3. Both control and SDC3-depleted myoblasts showed a robust response to serum stimulation, as expected since serum contains a large number of signaling molecules. However, the overall number of phosphopeptides that changed in SDC3-depleted myoblasts (Figure 2C and G, comparison #4 Δp-peptides = 1,416) was smaller than that in control myoblasts (Figure 2C and F, comparison #3 Δp-peptides = 1,614), consistent with the previous finding that in the presence of serum SDC3 depletion leads to fewer changes to the phosphoproteome than in the absence of serum (Figure 2D-E).

Lastly, we carried out a two-dimensional analysis to identify phosphosites that were sensitive to both serum stimulation and SDC3 presence/absence (Figure 2C, comparison #5 and Table S1). Not surprisingly, the number of phosphopeptides that significantly decreased as a consequence of both SDC3 depletion and serum stimulation (Figure 2C and H, comparison #5 and Table S1, Δp-peptides DOWN = 42) was greater than the number of phosphopeptides that increased (Figure 2C and H, comparison #5 and Table S1, Δp-peptides UP = 17).

To validate the phosphoproteomics data via a different approach, we obtained commercial antibodies that specifically recognize the phosphosites found sensitive to both serum and SDC3 (Table S1). Only for a few of these phosphosites could we find antibodies, including RPS6^S235/S236^ and RPS6^S240/S244^ (which were the top-most differentially abundant phosphopeptides sensitive to both serum and SDC3), IRS1^S307^ and IRS1^S302^. Western blotting analysis of phospho-RPS6 validated the phosphoproteomics data and showed that RPS6 phosphorylation both in serine 235/236 and in serine 240/244 were increased in serum-starved SDC3-depleted myoblasts compared with control myoblasts. Additionally, we validated the phosphoproteomic finding that RPS6 phosphorylation in both sites was enhanced by serum stimulation in control cells, but not in SDC3-depleted cells (Figure 2J-L). Of the IRS1 antibodies commercially available, only the antibody directed to IRS1^S302^ produced a detectable signal and showed no significant difference in response to either SDC3 depletion or serum stimulation (Figure 2M-O). However, it is well established that IRS1 is heavily regulated at the total protein level by many of its downstream effectors including S6 kinase, which also phosphorylates RPS6 (Harrington et al., 2005; Haruta et al., 2000; Shah and Hunter, 2006). Indeed, we observed that total IRS1 levels followed the same trend already described for phospho-RPS6 in response to both SDC3 depletion and serum stimulation (Figure 2M). All these observations further support the original hypothesis that SDC3 is somehow involved in repressing cell signaling in myogenesis.

### SDC3 regulates insulin signaling

Once established that SDC3 is heavily involved in the regulation of the myoblast phosphoproteome, we then asked which signaling pathways are regulated by SDC3. To address this question we mapped the proteins, and their associated phosphosites, regulated by both serum stimulation in control and SDC3-depleted myoblasts (Comparisons #3 and #4, Figure 2C) to canonical signaling pathways using Ingenuity Pathway Analysis (IPA). Given the large number of significantly enriched pathways returned by the IPA analysis (Table S2 and S3) and the relevance of tyrosine phosphorylation to this phosphoproteomic investigation (Figure 1H-I) we then restricted the functional enrichment analysis to tyrosine kinase-related signalling pathways (Table S4). Both control and SDC3-depleted myoblasts responded to serum by regulating similar signaling pathways, mostly related to growth factor and adhesion signaling (Figure 3A). In general, it appeared that the majority of tyrosine kinase-related pathways that were significantly enriched in response to serum were more enriched in SDC3-depleted cells compared to control cells (Figure 3A). However, nearly all of them were also less activated (smaller z-score) by serum in SDC3-depleted cells compared to control cells (Figure 3B). The most enriched pathway in SDC3-depleted myoblasts was *Integrin signaling* (Figure 3A). This supports our previous finding that a function of SDC3 in myoblasts and MuSCs is to regulate adhesion: primary *Sdc3^-/-^* MuSCs poorly adhere to myofibers and more easily than wild type MuSCs migrate off basal lamina-coated myofibers (Pisconti 2016). Consistently, we found that reduced adhesion to laminin is indeed a cell-autonomous property of myoblasts when SDC3 is depleted (Figure S2).

**Figure 3.**
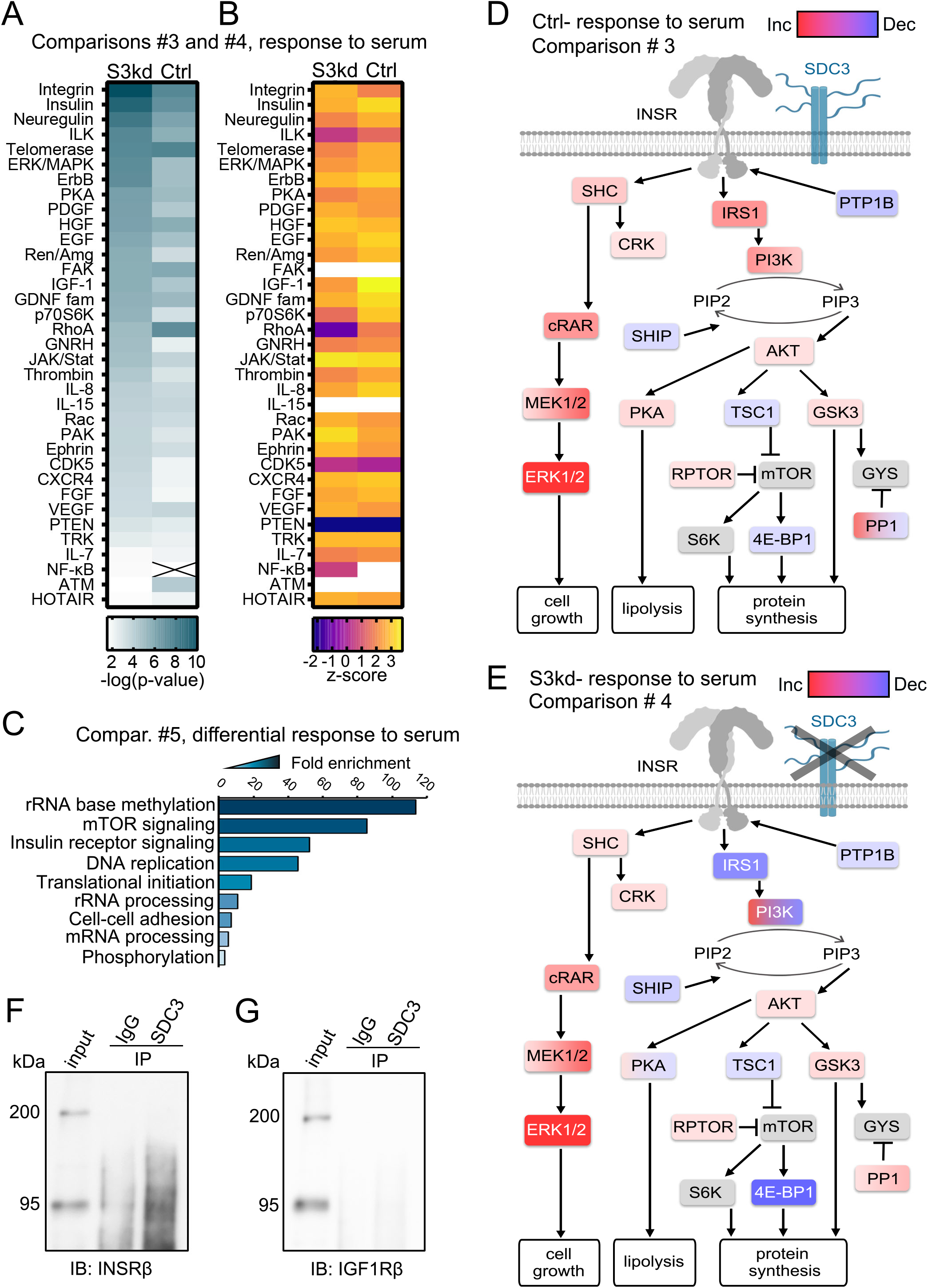
SDC3 coimmunoprecipitates with the insulin receptor and regulates proteins associated with insulin/AKT/mTOR signaling. **(A)** Phosphopeptides that were significantly up- or down-regulated when comparing the response to serum in control (Ctrl) and SDC3-depleted (S3^kd^) cells, were mapped to proteins with their associated phosphosite and submitted to Ingenuity Pathway Analysis (IPA). Pathway enrichment analysis was performed and the list of significantly enriched pathways was manually filtered to include only tyrosine kinase-related signaling pathways with a p-value < 0.05. An ‘X’ in the heatmap represents that the indicated pathway was not significantly enriched in the indicated cell line. Color scale: white = low enrichment, dark teal = high enrichment. **(B)** IPA’s z-score for signaling pathway enrichment analysis predicts whether a pathway is activated or deactivated: a negative value indicates deactivation whereas a positive value indicates activation. Color scale for pathways that were assigned an activation score: from orange to yellow = progressively more activated, from magenta to purple = progressively more deactivated. Where no prediction of activation level could be made the related cells are not colored. **(C)** Phosphopeptides identified as regulated by SDC3 and serum stimulation (comparison #5, Figure 2) were mapped to their corresponding proteins and submitted to DAVID 6.8 for gene ontology analysis. Biological Processes with a p<0.05 were considered significant, ordered by fold enrichment and plotted. **(D-E)** The insulin receptor pathway was interrogated in IPA for further analysis. Proteins and their phosphosites were mapped by IPA and colored according to increased (red) or decreased (blue) in abundance. Where multiple phosphosites were identified, the node was colored with shades of blue and/or red. Proteins shaded in gray are essential nodes of the pathways depicted for clarity but that were not found differentially phosphorylated. **(F-G)** C2C12 myoblasts were cultured in growth medium and detached using citric saline to avoid cleaving membrane complexes. Interacting plasma membrane proteins were fixed using 2% paraformaldehyde before lysing with an immunoprecipitation-friendly buffer. SDC3 was immunoprecipitated with its interactors using a rabbit anti-SDC3 antibody, while a normal rabbit IgG was used as control. Protein complexes were then treated to remove glycosaminoglycans, subjected to western blotting and probed for either (F) the insulin receptor (INSR) or (G) the insulin-like growth factor receptor (IGF1R). The IP’ed INSR appears as a smear due to variable levels of glycosylation it harbors (Hwang and Frost, 1999). IB = immunoblot. IP = immunoprecipitation. N = 3.

Next we interrogated the list of phosphoproteins that were differentially regulated by both SDC3 depletion and serum stimulation using the gene ontology (GO) analysis tool DAVID (Figure 3C). Interestingly, the significantly enriched (p-value < 0.05) *Biological Processes* to which these phosphoproteins mapped, were mostly related to protein translation, with *Insulin receptor* and *mTOR signaling* appearing at the top of the list as the most enriched processes. This was consistent with insulin signaling being the second most enriched signaling pathway in the response to serum (Figure 3A) and prompted us to further investigate the role of SDC3 in the regulation of insulin signaling in myogenesis.

Upon insulin stimulation, the insulin receptor (INSR) can signal via both the SHC/ERK pathway or via the PI3K/AKT/mTOR pathway, with the ERK pathway regulating mostly cell growth and the AKT/mTOR pathway regulating various metabolic functions such as glucose and lipid metabolism and protein translation (Boucher et al., 2014). Upon mapping to the INSR pathway the phosphoproteins that differentially responded to serum stimulation in SDC3-depleted and control myoblasts, it appeared that only the PI3K/AKT/mTOR branch of the pathway was differentially affected by the presence/absence of SDC3, while the SHC/ERK pathway was not (Figure 3D-E).

It is well-established that the IRS1/AKT/mTOR pathway is used by both insulin and insulin-like growth factors (IGFs), whose receptor insulin-like growth factor 1 receptor (IGF1R) shares a high degree of homology with the insulin receptor (INSR). Indeed, the INSR can be activated by IGF-1 and IGF-2, although it has greater affinity for its primary ligand insulin (Machackova et al., 2018). Therefore, to determine whether SDC3 selectively regulates the INSR or the IGF1R pathway, or both, we immunoprecipitated SDC3 from myoblasts and probed for both INSR and IGF1R. Despite the high homology between INSR and IGF1R, SDC3 immunoprecipitated only with the INSR but not IGF1R (Figure 3F-G). Given the higher affinity of INSR for insulin versus IGF1 and IGF2, we concluded that activation of insulin/mTOR signaling pathway mediated by serum in both SDC3-depleted and control myoblasts (Figure 3A) is likely due to insulin via INSR.

### SDC3 promotes insulin-induced proliferation but inhibits insulin-induced AKT phosphorylation in proliferating muscle progenitors

The role of insulin signaling in myogenesis is not completely understood. It appears to promote both myoblast proliferation and differentiation (Conejo et al., 2001; Grabiec et al., 2014; Grzelkowska-Kowalczyk et al., 2013; Mandel and Pearson, 1974). This is intriguing since myoblast proliferation and differentiation are two mutually exclusive biological processes and it is not clear through which mechanisms the same signaling molecule, insulin, is able to promote both. Thus, to test the hypothesis that SDC3 regulates insulin signaling in myogenesis, we cultured control and SDC3-depleted myoblasts (Figure 4A) as well as primary wild type and *Sdc3^-/-^* MuSC progeny (Figure 4B) in the presence/absence of increasing concentrations of insulin, measuring cell numbers before and after 24 hours treatment. Whilst control cell numbers increased in response to insulin treatment in a dose-dependent manner, the numbers of S3^kd^ myoblasts and *Sdc3^-/-^* MuSC progeny remained constant (Figure 4A-B), suggesting that SDC3 might be required for myoblast proliferation in response to insulin. To further test this, we measured the phosphorylation levels of AKT and ERK1/2 in control and SDC3-depleted myoblasts cultured in the presence/absence of insulin (Figure 4C-G). Neither ERK1 nor ERK2 phosphorylation were significantly increased by chronic insulin treatment of proliferating myoblasts (Figure 4C-E). In contrast, both control and SDC3-depleted myoblasts, cultured under proliferating conditions, responded to insulin by increasing the levels of AKT phosphorylation. However, SDC3-depleted cells, which fail to proliferate more in response to insulin, also showed greater activation of AKT in response to insulin than control cells (Figure 4F-G). Thus, hyperactivation of AKT signaling in response to insulin is associated with loss of proliferative response in SDC3-depleted myoblasts and primary *Sdc3^-/-^* MuSC progeny.

**Figure 4.**
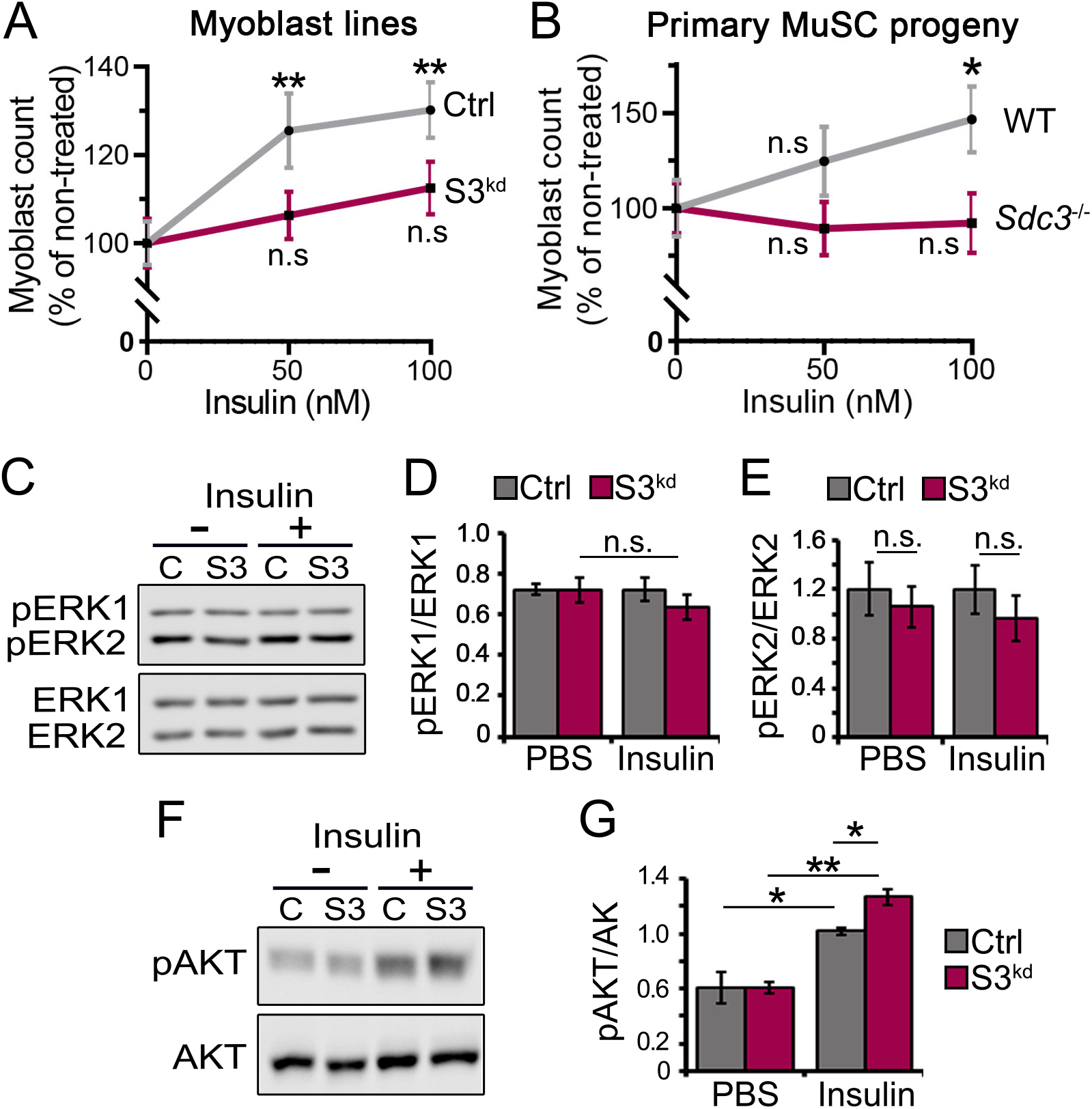
AKT phosphorylation induced by chronic insulin treatment is greater in proliferating S3^kd^ myoblasts compared to Ctrl myoblasts. **(A)** Control (Ctrl) and SDC3-depleted (S3^kd^) cells were cultured in growth medium for 24 hours supplemented with either PBS, 50 or 100 nM insulin. Cells were stained with DAPI to visualize nuclei and the number of cells per image was counted using a bespoke Fiji script. Data are plotted as a percentage of PBS treated cells. n.s. = non-significant, ** = p<0.01. Averages from 3 independent experiments, are plotted. Each biological replicate was plated in triplicate (3 biological replicates x 3 technical replicates per cell line) and 30 images per biological replicate were scored (total N□=□90 images per data-point). **(B)** Primary wild type (WT) and *Sdc3^-/-^* MuSC progeny were cultured and treated as in (A). Data expressed as a percentage of PBS treated cells. n.s. = non-significant * = p<0.05. Averages from 3 independent experiments are plotted, where 10 −15 images per biological replicate were scored (total N = minimum 30 images per data point). **(C)** Representative western blots of control (C) and SDC3-depleted (S3) myoblasts cultured in growth medium for 48 hours with either PBS or 100 nM insulin. Levels of phosphorylated-ERK1/2^T202/Y204^ (pERK1, pERK2) and total ERK1/2 (ERK1, ERK2) were measured. N = 3. **(D-E)** Quantification of western blotting images as those shown in (C) using ImageJ. n.s. = non-significant. Averages from 3 independent experiments, using a total of 3 biological replicates per cell line, are plotted. N = 3. **(F)** The same cell lysates from experiments shown in (C-E) were used to measure the levels of phosphorylated-AKT^S473^ (pAKT) and total AKT (AKT). N = 3. **(G)** Quantification of western blot images as those shown in (F). * = p<0.05, ** = p<0.01. Averages from 3 independent experiments, using a total of 3 biological replicates per cell line, are plotted. N = 3.

### SDC3 inhibits insulin-induced differentiation and insulin-induced AKT phosphorylation in differentiating muscle progenitors

Next, we tested the role of SDC3 in the regulation of insulin-induced myogenic differentiation in myoblast cell lines and primary MuSC progeny (Figure 5A). In the absence of additional insulin, S3^kd^ myoblasts differentiated and fused more vigorously than Ctrl myoblasts in response to serum lowering, as previously reported (Pisconti et al., 2010). Further insulin stimulation promoted differentiation and fusion in both cell lines, but Ctrl myoblasts responded to insulin to a greater extent than S3^kd^ myoblasts (Figure 5A-C). Similar results were obtained when primary MuSC progeny isolated from wild type and *Sdc3^-/-^* mice were tested (Figure 5D-F).

**Figure 5.**
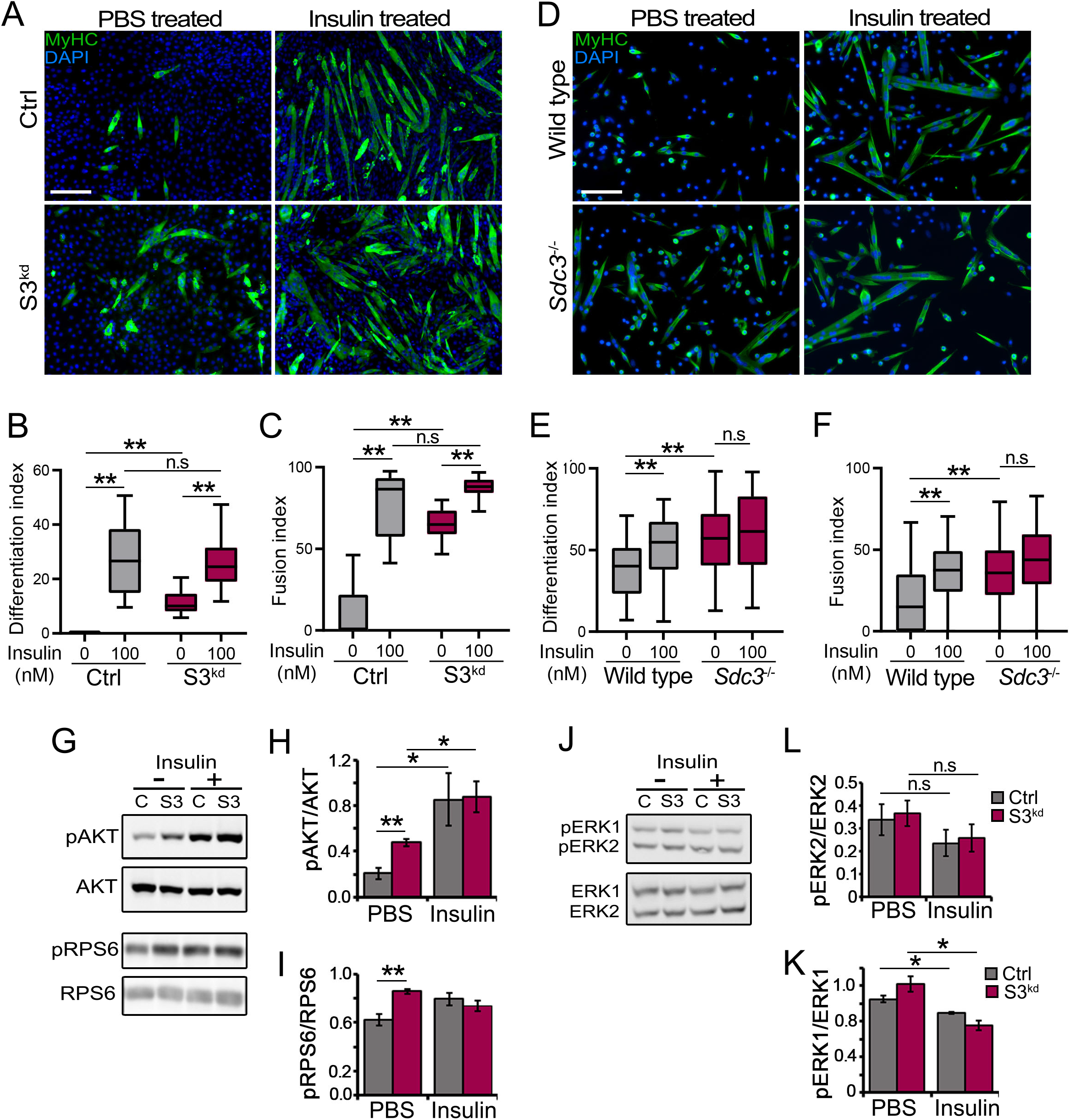
AKT/mTOR signaling is hyperactivated in differentiating SDC3-depleted myoblasts. **(A)** Representative immunofluorescent images of control (Ctrl) and SDC3-depleted (S3^kd^) myoblasts induced to differentiate for 3 days by reducing the serum concentration in the presence of PBS or 100 nM insulin. Cells were stained with DAPI (blue) to visualize nuclei and an anti-myosin heavy chain (MyHC, green) antibody (MF20 clone) to detect differentiated cells. Scale bar: 200 μm. **(B-C)** Quantification of fusion and differentiation index from cells treated as in (A). n.s. = non-significant. ** = p<0.01. Averages from 3 independent experiments are plotted. Each biological replicate was plated in triplicate (3 biological replicates x 3 technical replicates per cell line) and 10 images per biological replicate were scored (total N = 30). **(D)** Representative immunofluorescent images of primary wild type and *Sdc3^-/-^* MuSC progeny induced to differentiate for 3 days by reducing the serum concentration in the presence of PBS or 100 nM insulin. Cells were stained with DAPI (blue) to visualize nuclei and an anti-myosin heavy chain (MyHC, green) antibody (MF20 clone) to detect differentiated cells. Scale bar: 100 μm. N = 3. **(E-F)** Quantification of fusion and differentiation index from cells treated as in (D). n.s. = non-significant. ** = p<0.01. Averages from 3 independent experiments are plotted, where 15-20 images per biological replicate were scored (N = minimum 45 images per data point). **(G)** Control (C) and SDC3-depleted (S3) myoblasts were induced to differentiate for 3 days by reducing the serum concentration, in the presence of PBS or 100 nM insulin. Cells were lysed and subjected to western blotting to measure the levels of phosphorylated-AKT^S473^ (pAKT), AKT, phosphorylated-RPS6^S240/S244^ (pRPS6) or RPS6. N = 3, representative images of western blots are shown. **(H-J)** Quantification of western blotting images obtained as in (G) using ImageJ. * = p<0.05, ** = p<0.01. Averages of 3 independent experiments are plotted (N = 3). **(J)** The same cell lysates from experiments conducted as in (G) were used to measure the levels of phosphorylated-ERK1/2 (pERK1, pERK2) and total ERK1/2 (ERK1, ERK2). N = 3, representative images of western blots are shown. **(K-L)** Quantification of western blotting images as in (J). n.s. = non-significant. * = p<0.05. Averages of 3 independent experiments are plotted (N = 3).

When the levels of AKT and RPS6 phosphorylation were measured in differentiating myoblasts, as indicators of AKT/mTOR activation downstream of insulin, we observed a similar trend as shown by differentiation phenotypic markers: the absence of SDC3 led to an increase in basal AKT/mTOR activation in differentiating myoblasts, which was accompanied by a blunted response to insulin stimulation (Figure 5G-I). In contrast, ERK1/2 phosphorylation did not increase upon SDC3 loss in differentiating myoblasts, either under basal conditions or in response to insulin, although insulin stimulation caused a reduction in ERK1 phosphorylation in differentiating myoblasts, regardless of SDC3 expression (Figure 5J-K). This was not surprising since a reduction in ERK1/2 signaling is known to accompany myoblast differentiation (Flamini et al., 2018; Jones et al., 2001).

### SDC3 inhibits AKT phosphorylation in muscle progenitors thus preventing differentiation

The evidence accumulated so far indicates that SDC3 inhibits insulin-induced AKT phosphorylation, but not insulin-induced ERK1/2 phosphorylation, in both proliferating and differentiating myoblasts, but proliferation and differentiation are mutually exclusive processes in myoblasts. Importantly, SDC3 is downregulated during myoblast differentiation as previously reported (Fuentealba et al., 1999) and here further verified by us (Figure 6A). Thus, it is possible that SDC3 expression is the yet unknown “molecular switch” that converts myoblast response to insulin from pro-proliferative to pro-differentiative: when SDC3 expression is “on”, insulin signaling via AKT/mTOR is limited, thus inhibiting differentiation and promoting proliferation. When SDC3 expression is “off”, insulin signaling via AKT is amplified, thus promoting differentiation and the expense of proliferation. If this hypothesis is correct, then downregulation of the INSR in *Sdc3^-/-^* muscle progeny should restore differentiation back to wild type levels. To test this hypothesis, we transfected primary wild type and *Sdc3^-/-^* MuSC progeny with siRNAs targeting the INSR and induced them to differentiate by serum reduction (Figure 6B). Indeed, INSR knockdown in *Sdc3^-/-^* MuSC progeny inhibited differentiation and fusion and returned their values back to wild type levels (Figure 6C-D).

**Figure 6.**
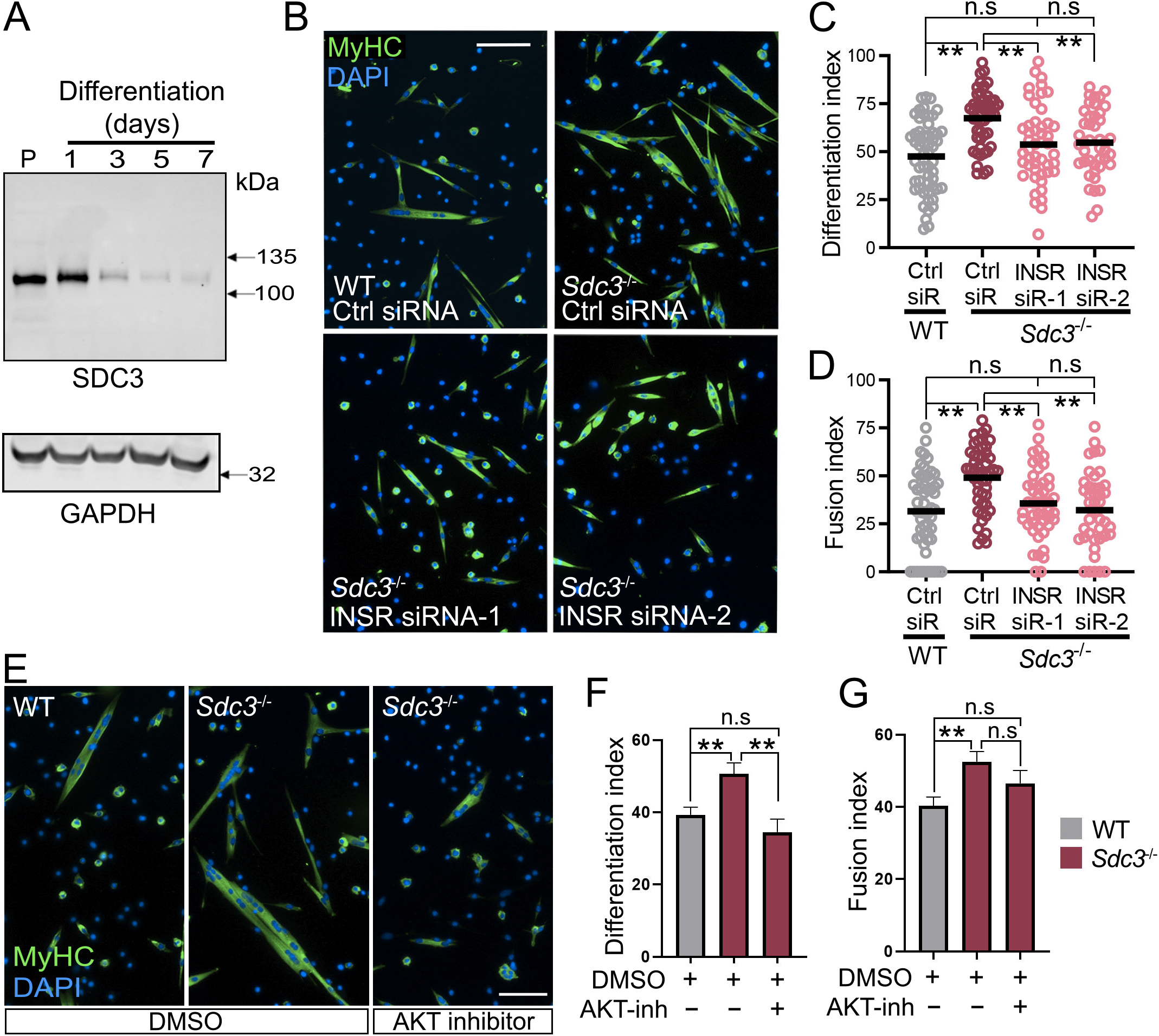
Knockdown of the insulin receptor and inhibition of AKT rescue the differentiation phenotype of *Sdc3^-/-^* MuSC progeny. **(A)** Protein levels of SDC3 in proliferating (P) and differentiating C2C12 myoblasts harvested at the indicated time points, were measured via western blotting. GAPDH was used as loading control. **(B)** Immunofluorescence images of primary wild type (WT) and *Sdc3^-/-^* MuSC progeny transfected with either 10 nM scrambled siRNA (Ctrl) or insulin receptor (INSR) siRNAs, then induced to differentiate for 40 hours by reducing serum concentration. Cells were stained with DAPI (blue) to visualize nuclei and with an anti-myosin heavy chain (MyHC, green) antibody (clone MF20) to detect differentiated cells. Scale bar = 100 μm. siR = siRNA. N = 3. **(C-D)** Quantification of fusion and differentiation index from primary (WT) and *Sdc3^-/-^* MuSC progeny treated as in (B). Values from individual images were plotted as individual dots, horizontal black lines are the averages calculated across 3 biological replicates per genotype, where 15-20 images per biological replicate were scored (N□=□minimum 45 images per data-point). n.s. = non-significant. ** = p<0.01. **(E)** Immunofluorescence images of primary wild type and *Sdc3^-/-^* MuSC progeny that were cultured in growth medium for four days leading to spontaneous differentiation. Cells were treated with DMSO or 1 μM AKT inhibitor (AKT inh) from four hours after plating. Cells were stained with DAPI (blue) to visualize nuclei and an anti-myosin heavy chain (MyHC, green) antibody (MF20 clone) to detect differentiated cells. Scale bar: 100 μm. N = 3. **(F-G)** Quantification of fusion and differentiation index from (E). Error bars represent S.E.M. ** = p< 0.01; n.s. = non-significant. Averages from 3 *Sdc3^-/-^* samples) and 4 (wild type samples) independent experiments are plotted, where 15-20 images per biological replicate were scored (N = minimum 45 images per data point).

Even in the presence of high serum concentrations, myoblasts that have proliferated and reached confluence will spontaneously begin to differentiate, which has been described as the community effect (Arnold et al., 2020; Cossu et al., 1995; Flamini et al., 2018). We hypothesized that, upon stochastic differentiation initiation, insulin contained in the serum is one of the signals which promotes spontaneous progression towards terminal differentiation and fusion. This effect would then be further amplified by SDC3 downregulation (Figure 6A) and consequent release of the SDC3-mediated break on insulin signal transduction via AKT/mTOR. If this hypothesis is correct, then: (i) the pro-differentiative effect of insulin during spontaneous differentiation in high serum should be exacerbated in *Sdc3^-/-^* MuSC progeny, where AKT/mTOR is not effectively inhibited during the proliferative phase; (ii) AKT inhibition should decrease *Sdc3^-/-^* MuSC progeny spontaneous differentiation to wild type levels. Indeed, when we cultured wild type and *Sdc3^-/-^* MuSC progeny under normal growth conditions (high serum) and allowed them to grow to confluence and then spontaneously differentiate (Figure 6E), SDC3 loss promoted both differentiation and fusion (Figure 6E). Furthermore, we confirmed that increased spontaneous differentiation in *Sdc3^-/-^* MuSC progeny could be rescued by AKT inhibition (Figure 6E-F). Importantly, AKT inhibition did not significantly affect myotube fusion (Figure 6G), suggesting that regulation of the insulin/INSR/AKT signaling pathway affects the earlier stages of differentiation, consistent with the dynamics of SDC3 downregulation upon differentiation initiation.

## DISCUSSION

We have used an unbiased phosphoproteomics approach to define the global landscape of signaling pathways regulated by SDC3 during myogenesis and identified insulin/AKT/mTOR as a key pathway regulated by SDC3 in MuSC progeny. This complements the recent finding that SDC3 regulates AKT in mesenchymal stem cells (Jones et al., 2020). Furthermore, we show that SDC3 is in a molecular complex with the INSR but not the IGF1R, suggesting this mechanism is specific to the INSR. Following an in depth investigation of SDC3-mediated regulation of insulin signal transduction, we discovered that SDC3 expression acts as a molecular switch allowing insulin to promote proliferation when SDC3 is expressed, in the early stages of myogenesis, while promoting differentiation when SDC3 is downregulated and eventually lost, in the later stages of myogenesis.

Several signaling molecules have been shown to promote both proliferation and differentiation of MuSC progeny, including insulin, however this is a biological conundrum as proliferation and differentiation are two mutually exclusive processes (Halevy et al., 1995; Ruijtenberg and van den Heuvel, 2016; Zhang et al., 1999). Here we show that in proliferating MuSC progeny SDC3 limits AKT activation in response to insulin, which in turn prevents premature differentiation (Figure 7A). As differentiation begins, SDC3 expression quickly decreases, releasing the inhibition on AKT and allowing differentiation to proceed (Figure 7B).

**Figure 7.**
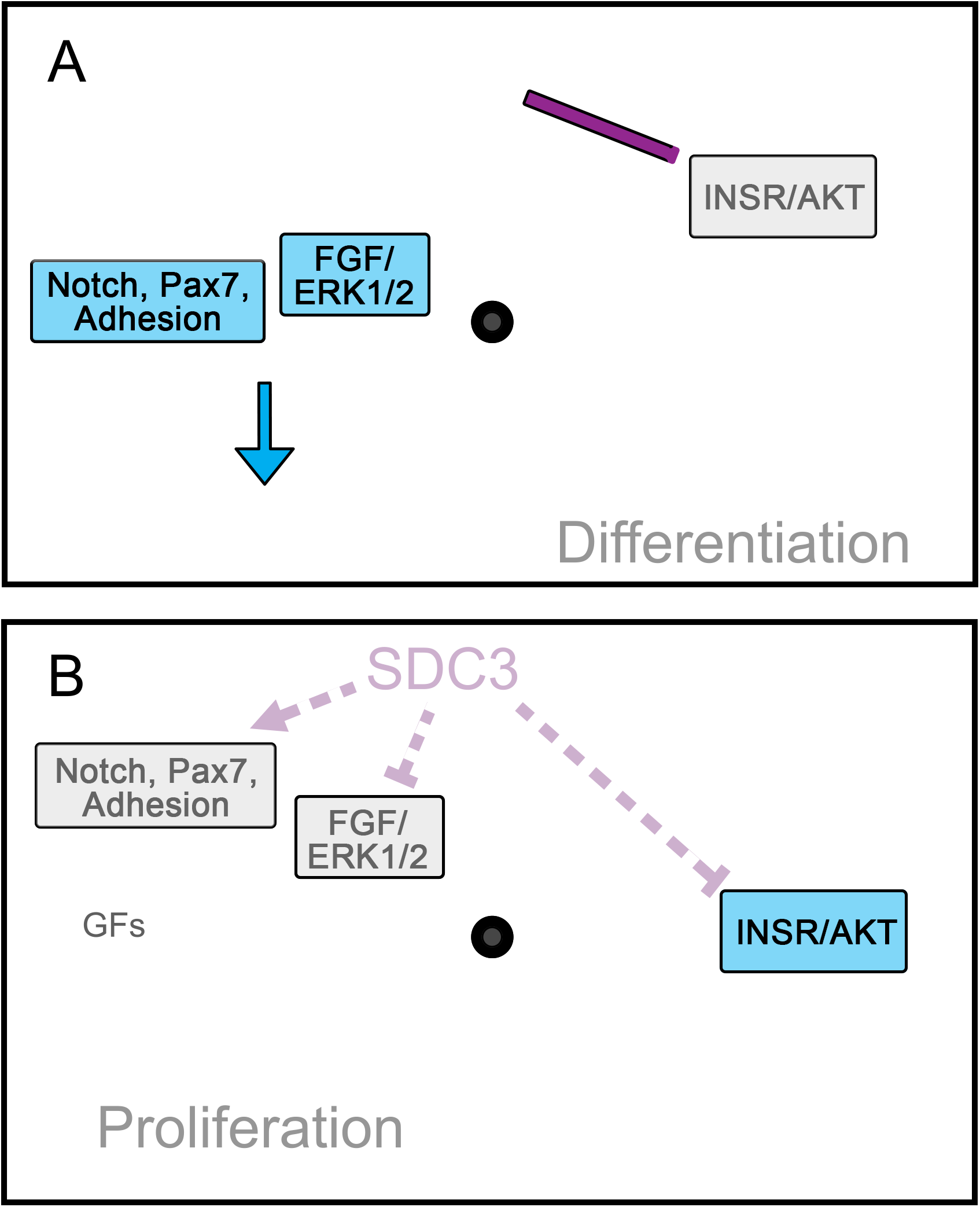
SDC3 is a timekeeper of myogenic differentiation. **(A)** During the early stages of myogenesis, SDC3 is expressed and supports proliferation by promoting Notch signaling, Pax7 expression and adhesion. At the same time SDC3 inhibits the insulin receptor (INSR)/AKT signaling pathway, which also supports proliferation by preventing premature differentiation. At this stage, growth factor (GF) signaling is also very active, further supporting proliferation, although a SDC3-mediated block on certain key GF pathways (e.g. FGF/ERK1/2 signaling pathway) poses a break to excessive GF signaling. **(B)** During the later stages of myogenesis, SDC3 is dramatically downregulated and its ability to promote proliferation is diminished whilst differentiation via Insulin/AKT is no longer inhibited. Additionally, the reduction in growth factor signaling further tips the balance towards terminal differentiation.

Inhibition of premature MuSC differentiation by SDC3 during the early stages of myogenesis is an essential strategy to ensure that the MuSC progeny pool expands enough to yield the numbers of progeny necessary for a complete regeneration (Figure 7). Indeed, as shown by us and others, SDC3 fulfils this timekeeper role also via regulation of other signaling pathways. We previously showed that SDC3 promotes MuSC progeny expansion by enhancing cleavage and activation of Notch1, which in turn inhibits differentiation and promotes cell cycle progression (Buas and Kadesch, 2010; Pisconti et al., 2010). Moreover, SDC3 is required for Pax7 expression and maintenance of an uncommitted state (Pisconti et al., 2016), which is an essential mechanism in the prevention of differentiation (Olguín and Olwin, 2004; Olguin et al., 2007), and promotes MuSC adhesion to myofibers, which is required for quiescence maintenance (Goel et al., 2017).

An observation apparently at odds with the pro-differentiative effect of SDC3 downregulation upon differentiation initiation is that SDC3 appears to inhibit FGF2 and HGF signaling via ERK1/2 (Cornelison et al. (2004) and Figure 1C. Therefore, loss of SDC3 expression during the later stages of myogenesis should lead to ERK1/2 hyperactivation, but this would inhibit differentiation (Knight and Kothary, 2011), while we observe enhanced differentiation when SDC3 is absent or heavily depleted. In reality, expression of ERK1/2 is also rapidly downregulated with the onset of differentiation (Flamini et al., 2018), therefore positive regulation of ERK1/2 activity in the absence of SDC3 is only relevant during the early, proliferative stages of myogenesis, but not during the late, differentiative stages of myogenesis. Moreover, the overall heparan sulfate complement of MuSC progeny changes upon differentiation onset, becoming inhibitory for FGF2 signaling via ERK1/2 (Ghadiali et al., 2016). In fact, it is possible that SDC3-mediated inhibition of ERK1/2 in response to growth factors is the reason why regenerated myofibers are larger in *Sdc3^-/-^* mice compared to wild type controls. In the absence of SDC3, loss of proliferative capacity due to impairments in adhesion, Pax7 expression and Notch signaling and to enhanced INSR-mediated differentiation, is partly counteracted by enhanced ERK1/2 activation in response to growth factors, effectively preserving expansion of the MuSC progeny pool. Moreover, impaired Notch signaling and reduced adhesion to myofibers lead to loss of MuSC quiescence in *Sdc3^-/-^* muscles, which contributes to further expand the pool of differentiation-competent MuSC progeny, ultimately leading to generation of larger myofibers (Cornelison et al., 2004; Pisconti et al., 2016; Pisconti et al., 2010). In this context, it is also possible that SDC3-mediated inhibition of ERK1/2 activation by growth factors has evolved as sort of “break” preventing excessive MuSC proliferation and eventual tumor formation (Rees et al., 2006; Taylor et al., 2009), which places SDC3 as a balancer at the center of an intricate signaling network (Figure 7A).

Integrin signaling was amongst the top signaling pathways regulated by SDC3. Integrin-mediated cell adhesion may also influence insulin-mediated differentiation since integrins bind to the INSR and IRS1 upon insulin stimulation to mediate insulin signaling and promote proliferation (Vuori and Ruoslahti, 1994). Additionally, insulin induces dephosphorylation of focal adhesion kinase (FAK) in adherent cells and FAK/IRS1 have been shown to interact, which strongly suggests co-operation of insulin and integrin signaling (El Annabi et al., 2001). Loss of SDC3 perturbs MuSC adhesion (Pisconti et al., 2016) and enhances differentiation. Therefore, SDC3 might regulate an insulin/integrin signaling cross-talk as syndecans cooperate with integrins to mediate cell adhesion to the ECM (Beauvais and Rapraeger, 2003; Whiteford et al., 2007). A hypothetical model may include an integrin/INSR/SDC3 interaction in MuSC progeny that promotes proliferation in response to insulin; when SDC3 is lost this interaction is impaired and differentiation is promoted via AKT. The integrin/INSR/SDC3 complex would then prevent differentiation by inhibiting IRS1/AKT signaling, which is supported by the significant changes in the levels of total and phosphorylated IRS1 reported in this paper.

Diabetic myopathy is the reduction in muscle mass and function associated with type 1 and 2 diabetes mellitus. Skeletal muscle is heavily involved in regulating blood glucose, and in diabetic patients the total muscle mass is reduced (Tajiri et al., 2010). MuSCs differentiation is impaired in diabetic mice, leading to a lack of muscle regeneration in response to injury (D′Souza et al., 2013). Additionally, MuSC proliferative capacity is reduced in a rat model of metabolic syndrome, accompanied by alterations to AKT signaling and myogenic regulatory factor expression (Peterson et al., 2008). Changes in SDC3 expression have been linked to metabolic syndrome, regulation of energy balance and obesity, all risk factors predisposing to development of type 2 diabetes mellitus (Chang et al., 2018; Strader et al., 2004). Moreover, it has been shown that fetal hyperglycemia caused by gestational diabetes of the mother causes insulin resistance in developing skeletal muscle (Kua et al., 2019). Further work to investigate the role of SDC3/INSR/AKT signaling in a diabetic model would be advantageous to elucidate any protective effects from targeting this pathway on diabetic myopathy as well as on regulation of glucose levels.

In conclusion, our work, and that of others, establish SDC3 as an important regulator of myogenesis in charge of coordinating the delicate balance between multiple signaling pathways and biological processes. This suggests that its pharmacologic targeting might result beneficial in the context of adult muscle regeneration, developmental and obesity-related disorders.

## Acknowledgments

The authors wish to thank the members of the Centre for Proteome Research at the University of Liverpool, especially Prof Claire Eyers and Dr Philip Brownridge. We thank Prof Dave Fernig for kindly donating recombinant FGF2 and Prof Alan Rapraeger for the rabbit anti-SDC3 antibody. This work was funded by the Biotechnology and Biological Sciences Research Council (BBSRC) in the form of DTP funding to F.K.J. and A.P.*, a Wellcome Trust ISSF fellowship to A.P.*.

* = Addolorata Pisconti

## Author Contributions

Conceptualization, A.P* and F.K.J. Methodology; A.P*, F.K.J, A.J and A.P; Software, A.J and A.P; Formal Analysis, A.P*, F.K.J, A.J and A.P; Investigation, A.P* and F.K.J; Resources, A.P* and A.J; Writing – Original Draft, A.P* and F.K.J; Writing – Review & Editing, A.P*, F.K.J, A.J and A.P; Visualization, A.P* and F.K.J; Supervision, A.P* and A.J; Project Administration, A.P* and F.K.J; Funding Acquisition, A.P*.

* = Addolorata Pisconti

## Materials and methods

A full list of reagents, tools and software used, with their unique identifiers, are available in Supplemental Table 5.

### Data and Code Availability

The mass spectrometry proteomics data have been deposited to the ProteomeXchange Consortium via the PRIDE (Perez-Riverol et al., 2019) partner repository with the dataset identifier PXD020123.

The Bayesian Markov chain Monte Carlo (MCMC) code is available from https://github.com/PGB-LIV/JonesSDC3PhosphoproteomicsPaper

The Fiji script used to count cell nuclei and myoblast area is available from https://github.com/Piscontilab/Fiji-script

### Animals

Wild type (C57Bl/6) and age and sex-matched *Sdc3^-/-^* mice (Reizes et al., 2001) were housed in a pathogen-free facility at the University of Liverpool in accordance with the Animals (Scientific Procedures) Act 1986 and the EU Directive 2010/63/EU, after the local ethical review and approval by Liverpool University’s Animal Welfare and Ethical Review Body.

### C2C12 cell cultures

Routine culture of parental C2C12, Ctrl and S3^kd^ myoblasts was performed under humidified atmosphere of 5% CO_2_ and atmospheric O_2_ at 37 °C. Growth medium was composed of DMEM (D6429 Sigma-Aldrich), 10% foetal bovine serum (Gibco), 1% penicillin and streptomycin (Invitrogen). Cells were passaged every two days to ensure confluence was kept between 40-70%. Parental C2C12 cells were obtained from ATCC and passages did not exceed 30. Cells were detached for passaging using 0.5% trypsin-EDTA solution (Sigma, T3924). For routine differentiation of parental C2C12, Ctrl and S3^kd^ myoblasts, 90% confluent cultures were washed twice with DMEM before being cultured in differentiation medium (DMEM, 2% horse serum, 1% pen/strep) for various lengths of time.

### Primary muscle stem cell-derived myoblast cultures

Primary muscle stem cell-derived myoblasts (henceforth referred to as MuSC progeny) were obtained from hindlimbs of 8-14 week old male mice (wild type and age-matched *Sdc3^-/-^*) as described previously (Flamini et al., 2018; Ghadiali et al., 2016). Growth medium consisted of F12 + 0.4 mM CaCl2 (F12C), 15% horse serum (HyClone), 1% penicillin/streptomycin, 2 nM recombinant FGF2 (kindly donated by Dr David Fernig, University of Liverpool). MuSC progeny were passaged once, two days after initial isolation and plating, using 40 units/mL of collagenase type I in phosphate buffered saline (PBS). Differentiation medium consisted of F12C, 3% horse serum, 1% penicillin/streptomycin.

### Generation of Ctrl and S3^kd^ myoblasts

SDC3 was stably knocked down in C2C12 myoblasts by transfecting cells with a plasmid containing an shRNA sequence that targets SDC3 (Sigma, Mission # TRCN0000071990) to generate the SDC3-depleted (S3^kd^) cell line, or the empty backbone vector (pLKO-1, Sigma Mission) to generate the control (Ctrl) cell line. C2C12 myoblasts at 80% confluence were transfected using Lipofectamine 2000 then passaged for 3 times in the presence of 2 μg/mL puromycin to select for positively transfected cells. No clonal selection was employed since C2C12 myoblasts are heterogenous. Western blotting was used to confirm knockdown efficiency, which was ≥ 95%.

### Western Blotting

Protein extracts were obtained by lysing cells in RIPA buffer (50 mM Tris-HCl, pH 7.4, 1% NP-40, 0.25% sodium-deoxycholate, 150 mM NaCl, and 1 mM EDTA) supplemented with a protease inhibitor cocktail (Complete, Roche) and phosphatase inhibitors (2 mM activated Na_3_VO_4_, 2 mM NaF and 1x PhosSTOP (Roche)). Soluble proteins were quantified using the Pierce BCA Protein Assay Kit, following the manufacturer’s protocol. For detection of all proteins indicated except SDC3, an equal amount of protein sample (between 10-20 μg) was separated by SDS-PAGE and then transferred onto a nitrocellulose membrane (Hybond, GE Healthcare) for two hours at 250 mA on ice. Membranes were blocked with a solution of 5% non-fat milk in Tris Buffered Saline Tween20 (TBST). Primary antibodies (see Supplemental Table 5) were diluted in either 5% w/v BSA in TBST or 5% milk in TBST, depending on the supplier’s recommendation, added to membranes and incubated over night at 4 °C. Membranes were washed in TBST and incubated with a horseradish peroxidase (HRP)-conjugated secondary antibody for one hour at room temperature. After washing with TBST, membranes were visualised using chemiluminescence (Clarity™ ECL, Biorad) and imaged on an ImageQuant-Las4000 (GE healthcare) gel doc system or Amersham Imager 680. Band intensity was analysed using the ‘Analyze Gel’ function in ImageJ.

To detect SDC3, cells were lysed as described above and equal amounts of Ctrl and S3^kd^ total protein were precipitated with 2.5x volume of methanol overnight at −20 °C. Precipitates were pelleted by centrifugation at 13,000 x g at 4 °C then washed once with −20 °C acetone and centrifuged again at 4 °C to collect precipitates. After discarding the supernatant, pellets were re-suspended in heparinase buffer (100 mM sodium acetate, 0.1 mM calcium acetate and 1 mM magnesium chloride) and digested with 0.25 mU heparinase III (Ibex) and 0.5 mU chondroitinase-ABC (Sigma-Aldrich) for 2 hours at 37 °C, before addition of another 0.25 mU heparinase III for an additional two hours. Reactions were quenched by addition of 5X SDS-sample buffer and heated to 95°C for 10 min before 50 μg of protein samples were loaded onto SDS-PAGE. From this point western blotting were carried out as described above.

### Stimulation of Ctrl and S3^kd^ myoblasts with FGF2 or serum

S3^kd^ and Ctrl cells were plated at a density of 45,000 cells per 10 cm plates and cultured for two days. For serum-starvation, cells were washed twice with DMEM and then incubated for 5 hours in DMEM + 2 μg/mL puromycin. After serum-starvation myoblasts were either lysed as described above in the western blotting section or treated with either 10% FBS or 2 nM FGF2 and then lysed at various time points as described in the western blotting section.

### Immunostaining, microscopy and quantification

Culture medium was aspirated, and cells fixed with 4% paraformaldehyde (Sigma) in PBS (pH 7.4) solution for 10 minutes at room temperature. Cells were permeabilised with PBS + 0.2% TritonX100 for 10 minutes at room temperature, then washed once with PBS + 0.2% TritonX100 and twice with PBS prior to immunostaining. For detection of myosin heavy chain (MyHC), the primary anti-MyHC antibody (DSHB, MF20 clone) was diluted at 1:150 in PBS + 1% horse serum before being incubated with cells overnight at 4°C. Next, cells were washed as above before a secondary antibody (anti-mouse conjugated to AlexaFluor488) was diluted at 1:500 in PBS + 5% horse serum and incubated with cells for 1 hour at room temperature. A solution of 2 μg/mL DAPI (Life Technologies) in PBS was used to stain DNA. Cells were then washed as above and stored in the dark at 4°C until imaging.

Images were taken using an epifluorescence microscope (EVOS-FL Life Technologies) with the same exposure and gain settings in each experiment.

Quantification of immunostaining was performed by taking a minimum of 20 random images per experiment, with three technical replicates and three biological replicates unless otherwise stated. A bespoke script written for Fiji was used to count DAPI+ nuclei and myotube area as described previously (Arecco et al., 2016). Differentiation was calculated as the number of nuclei in MyHC positive cells divided by the total number of nuclei x100. Fusion index was calculated as the number nuclei contained in MyHC positives myotubes with at least two nuclei, divided by the total number of nuclei in all MyHC positive cells x100.

### Phosphoproteomics sample preparation and LC-MS/MS

Lysates were prepared as previously described (Jones et al., 2019). Ctrl and S3^kd^ myoblasts were cultured in growth medium for two days and for each condition four biological replicates were used. Myoblasts were serum-starved for 5 hours in DMEM + 2 μg/mL puromycin then stimulated with FBS for 10 min. Cells were lysed in a mass spectrometry compatible buffer (50 mM ammonium bicarbonate, 0.25% RapiGest^™^ (Waters, UK), 2 mM NaF, 2 mM Na_3_VO_4_ and 1x PhosSTOP). Lysates were rotated end-over-end for 30 minutes at 4°C, then sonicated with a Vibracell ultrasonic processor (Sonics, Newtown, USA) on ice for 10 seconds with a tapered microtip. Lysates were then snap-frozen in liquid nitrogen and submitted to the Centre for Proteome Research (CPR), University of Liverpool, for LC-MS/MS analysis. Briefly, 200 μg of protein sample were digested with 0.2 μg/μL trypsin overnight at 37 °C. RapiGest™ was hydrolysed by the addition of trifluoroacetic acid (TFA), for 2 hours at 60 °C, followed by centrifugation at 17,000 x g for 30 minutes at 4 °C. Supernatants were concentrated using a speedvac (Univapo-150 ECH) at a fixed speed of 1,250 rpm, 30 °C for 40 minutes and resuspended in 150 μl of 1% TFA, then desalted on macro spin columns (Harvard Macro Spin Columns). Samples were enriched for phosphopeptides using Titansphere PhosTiO columns (200 μL 5010-21312, Hichrom) prior to being solubilised in 20 μL 1% TFA, and centrifuged at 17,000 *x g* for 15 minutes. 10 μL of supernatant were transferred to total recovery vials for LC-MS/MS analysis.

LC-MS/MS was performed at the Centre for Proteome Research, University of Liverpool, UK. Briefly, 4 μL of the phospho-enriched sample were analysed using an Ultimate 3000 RSLC™ nano system (Thermo Scientific, Hemel Hempstead) coupled to a QExactive™ mass spectrometer (Thermo Scientific). The sample was loaded onto the trapping column (Thermo Scientific, PepMap100, C18, 300 μm x 5 mm), for 7 minutes at a flow rate of 9 μL/min with 0.1% TFA and 2% acetonitrile. The sample was resolved on the analytical column (Easy-Spray C18 75 μm x 500 mm 2μm column) using a gradient of 97% buffer A (0.1% formic acid in water) and 3% buffer B (99.9% acetonitrile, 0.1% formic acid) to 60% buffer A and 40% buffer B (all v/v) over 90 minutes at a flow rate of 300 nL/min. The data-dependent program used for data acquisition consisted of a 70,000 resolution full-scan MS scan (automatic gain control (AGC) set to 1 x106 ions with a maximum fill time of 250 ms) and the 10 most abundant peaks were selected for MS/MS using a 35,000 resolution scan (AGC set to 1 x 105 ions with a maximum fill time of 100 ms) with an ion selection window of 2 m/z. To avoid repeated selection of peptides for MS/MS, the program used a 20 second dynamic exclusion window.

### Peptide identification and label-free quantification

Peptides were identified using PEAKS Studio 8 searched against the Uniprot *Mus musculus* proteome database (Database version: October 2016). A fixed carbamidomethyl modification for cysteine and variable modifications of oxidation for methionine were specified. A variable modification for phosphorylation of serine, threonine and tyrosine were specified. A precursor mass tolerance of 10 ppm and a fragment ion mass tolerance of 0.05 Da were applied. The results were then filtered to obtain a peptide false discovery rate of 1%. The localisation probability of all putative phosphorylation sites were determined by PEAKS and reported as the Ascore. No filtering of peptides by Ascore was performed.

Chromatogram alignment and label free quantification was performed using Progenesis QI (Version 2.0). Bayesian Markov chain Monte Carlo (MCMC) simulation was performed using the Stan (DOI 10.18637/jss.v076.i01) programming language to fit a Poisson regression model with a log-link for each peptide in each comparison. In cases where a peptide was unobserved in a condition, an informative prior was used to impute a range of potential missing values for that condition. For each condition, a log-linear regression of the peptide abundances against the number of observed data points was used to predict the distribution of abundances of peptides with no observed values. This predicted distribution was then used as an informative prior for the abundance of missing values.

For comparisons #1 - #4, the inferred posterior distributions of the mean log-fold-changes were then used to perform one-sided significance tests to determine the probabilities that the mean fold-change was less than −1.5 or greater than +1.5 for each peptide in each comparison. The complement of the greater of these two probabilities was taken as the quantitative posterior error probability (PEP) for each peptide in each comparison. For comparison #5, the posterior samples of mean log-fold-change in comparisons #3 and #4 were used to determine the posterior distribution of the difference in log-fold-change between comparison #3 and #4 for each peptide and quantitative PEPs calculated as above. Significant sets of peptides for each of the five comparisons were determined by calculating the largest set where the mean PEP <0.05.

### Pathway enrichment analysis

Significantly regulated phosphopeptides and their corresponding Uniprot protein ID, natural log fold change and phosphosite were submitted to Ingenuity Pathway Analysis (IPA- Version 2.3, Qiagen) for canonical pathway enrichment using the phosphoproteome analysis tool. Pathways with an enrichment score of p <0.05 were considered statistically significant. IPA also returns a z-score which gives a prediction of whether pathways detected lead to activation (positive z-score) or deactivation (negative z-score) of that signaling pathway, depending on the proteins and phosphosites that are mapped to that pathway.

For Gene Ontology enrichment analysis, DAVID 6.8 (Huang da et al., 2009a, b) was used with significantly regulated phosphoproteins. The background dataset consisted of all the phosphoproteins identified in the phosphoproteomics experiment. Enriched terms with a p-value of less than 0.05 were considered statistically significant.

### Co-immunoprecipitation of SDC3 interactors

Immunoprecipitation of SDC3 interactors was performed by fixation of cell lysate as previously described with small modifications (Klockenbusch and Kast, 2010). Briefly, C2C12 myoblasts were cultured in normal growth promoting conditions and then gently detached using citric saline (135 mM potassium chloride and 15 mM sodium citrate, no pH adjustment). Cells were pelleted and resuspended to 10^7^ cells/mL in 2% paraformaldehyde/PBS solution and incubated for 10 minutes. Cells were pelleted and washed in 0.5 mL ice-cold 1.25 M glycine/PBS solution twice. Cells were then lysed in 1 mL per 1×10 cells with a lysis buffer optimized for immunoprecipitation of heparan sulfate proteoglycans (1 X PBS, 1% Triton X100, 0.1% SDS), for 30 min at 4°C with rotation. After 30 minutes cells were further lysed by passing through a 27G needle 10 times. Lysates were then centrifuged at 17,000 x g at 4 °C for 30 min, soluble extracts were snap-frozen and stored at −80°C.

Protein concentration was determined using the Pierce BCA Protein Assay Kit, following the manufacturer’s protocol. 1mg of lysate was diluted in PBS to a volume of 1 mL with the addition of either anti-SDC3 antibody (kindly donated by Dr Alan Rapraeger, University of Wisconsin at Madison, and previously validated by us (Pisconti et al., 2010) or normal-rabbit IgG. Antibodies and cell lysate were incubated overnight at 4 °C with gentle rotation. The next day, immunocomplexes were isolated using protein A agarose beads as directed by the manufacturer’s protocol (Pierce, Thermo Fisher). Whilst immunocomplexes were still attached to the beads, glycosaminoglycan chains were removed using heparinase III and chondroitinase ABC as described in the western blotting section. After digestion, immunocomplexes were eluted from the beads by boiling the samples in 2X SDS buffer. Equal volumes of eluted samples were then resolved on an 8% SDS-PAGE and western blotting performed as described above.

### Isolation of primary muscle stem cell-derived myoblasts

Hindlimb muscles were dissected and minced before digestion with 400 units/mL collagenase type I (Worthington Biochemical Corporation) in F12C for 1.5 hours at 37 °C with mixing every 15 minutes. Cell pellets were collected and resuspended in primary growth medium before being filtered through a 40 μm cell strainer. Cell pellets were resuspended again and plated onto gelatin-coated dishes under normal culture conditions for 1 hour. Non-adherent cells and growth medium were collected and then re-plated onto new gelatin-coated plates for two hours. For the final plating, non-adherent cells were collected, centrifuged, re-suspended in fresh growth and plated onto fresh gelatin-coated plates for routine culturing.

### Stimulation of Ctrl and S3^kd^ and control cells with insulin

For proliferation experiments, Ctrl and S3^kd^ cells were seeded at an equal density in growth medium supplemented with 50 or 100 nM insulin (I0516, Sigma Aldrich) as indicated in figures, or PBS as vehicle control, for 24 hours. During insulin treatment, primary MuSC progeny were maintained without FGF2. After 24 hours, both primary cells and cell lines were fixed and stained with DAPI as described in the immunostaining, microscopy and quantification section.

To measure differentiation and fusion in response to insulin, cells were grown to ~90% confluence before switching them to differentiation medium. All cells were induced to differentiate for three days in the presence of 100 nM insulin or PBS. Cells were then fixed and stained with DAPI and anti-MyHC antibody as described in the immunostaining, microscopy and quantification section.

To measure the levels of pAKT, AKT, pRPS6, RPS6, ERK1/2 and ERK in proliferating myoblasts, Ctrl and S3^kd^ cells were cultured for two days in growth medium supplemented with 100 nM insulin or PBS. To measure the same protein levels in differentiating cells, Ctrl and S3^kd^ cells were grown to ~90% confluence and then switched to differentiation medium containing either 100 nM insulin or PBS. Cells were differentiated for three days. After the allotted time had passed, cells were lysed and processed for western blotting as described in the relevant section.

### Transfection of insulin receptor siRNA in primary myoblasts

For transient knockdown of INSR, primary MuSC progeny were seeded at 8,000 cells/well in 12-multiwell plates in growth medium. For induced differentiation experiments, cells were cultured in growth medium for two days before medium was switched to transfection medium (Opti-MEM + 15% horse serum + 2 nM FGF2), whilst lipofectamine 2000 plus 10 nM INSR siRNAs or scrambled siRNA (sequences in Key Resources Table) were prepared. Lipofectamine 2000 was used as recommended by the manufacturer. Cells were incubated in lipofectamine2000 + siRNAs for five hours and then complexes removed and the culture medium switched to differentiation medium. Cells were left to differentiate for 40 hours before fixation.

For spontaneous differentiation experiments, cells were cultured in growth medium for two days before being switched to transfection medium whilst lipofectamine + siRNAs were being prepared as described above. After five hours the transfection medium was removed and replaced with growth medium minus FGF2 for 48 hours. Cells were fixed and stained as described in the immunostaining, microscopy and quantification section.

### Inhibition of AKT in primary MuSC progeny

Primary MuSC progeny were seeded at a density of 7,000 cells/well in 12-multiwell plates in growth medium containing FGF2 and allowed four hours to adhere to the gelatin-coated plates before beginning treatment with an AKT inhibitor (AZD5363, SelleckChem). After culturing in growth medium for 2-3 days, until confluent, FGF2 was removed (by replacing the medium with fresh growth medium that did not contain FGF2) and cultured for an additional two days still in the presence of the AKT inhibitor. A volume of DMSO equal to the volume of AKT inhibitor, was added to vehicle control wells. Cells were then fixed and stained with DAPI and anti-MyHC antibody as described above.

### Adhesion assay

Ctrl and S3^kd^ myoblasts were detached by trypsin, centrifuged, re-suspended in C2C12 growth medium and counted using a hemocytometer. An equal number of Ctrl and S3^kd^ cells were added to two separate tubes and centrifuged to remove growth medium before being washed twice with either growth medium or serum-free medium (DMEM + 1% penicillin/streptomycin). After the second wash, both sets of cells were centrifuged and re-suspended at 7,000 cells/mL in either DMEM alone or complete growth medium (DMEM + serum). Cells were incubated for 1 hour at 37 C, 5% CO_2_ before washed once with 1x PBS to remove non-adherent cells, then fixed in 4% PFA for 10 minutes at room temperature and then stained with DAPI to visualise cell nuclei before the number of cells were counted using the bespoke script as described above.

### Quantification and statistical analysis

All statistical analysis for biochemistry and immunofluorescence experiments were performed in GraphPad Prism 8. Each experiment contained at least three biological replicates and multiple technical replicates. Data were checked for normal distribution and statistical significance was calculated using the Student’s t-test or one-way ANOVA followed by Tukey post-test. Data were considered significant if p < 0.05. Data were plotted as average ± SEM (standard error of the mean) where error bars are shown.

## Supplemental Information Titles and Legends

Table S1. List of phosphopeptides with significant changes in abundance with both serum stimulation and SDC3 presence/absence, related to Figure 2.

Table S2. List of significantly enriched pathways returned by the IPA analysis of serum stimulated Ctrl cells, related to Figure 3

Table S3. List of significantly enriched pathways returned by the IPA analysis of serum stimulated S3^kd^ cells, related to Figure 3

Table S4. List of tyrosine kinase-related signalling pathways, related to Figure 3.

Table S5. List of reagents and resources used.

## References

Arecco, N., Clarke, C.J., Jones, F.K., Simpson, D.M., Mason, D., Beynon, R.J., and Pisconti, A. (2016). Elastase levels and activity are increased in dystrophic muscle and impair myoblast cell survival, proliferation and differentiation. Scientific reports 6, 24708.

Arnold, L.L., Cecchini, A., Stark, D.A., Ihnat, J., Craigg, R.N., Carter, A., Zino, S., and Cornelison, D. (2020). EphA7 promotes myogenic differentiation via cell-cell contact. Elife 9.

Beauvais, D.M., and Rapraeger, A.C. (2003). Syndecan-1-mediated cell spreading requires signaling by alphavbeta3 integrins in human breast carcinoma cells. Exp Cell Res 286, 219–232.

Boucher, J., Kleinridders, A., and Kahn, C.R. (2014). Insulin receptor signaling in normal and insulin-resistant states. Cold Spring Harb Perspect Biol 6.

Buas, M.F., and Kadesch, T. (2010). Regulation of skeletal myogenesis by Notch. Exp Cell Res 376, 3028–3033.

Casar, J.C., Cabello-Verrugio, C., Olguin, H., Aldunate, R., Inestrosa, N.C., and Brandan, E. (2004). Heparan sulfate proteoglycans are increased during skeletal muscle regeneration: requirement of syndecan-3 for successful fiber formation. J Cell Sci 117, 73–84.

Chang, B.C., Hwang, L.C., and Huang, W.H. (2018). Positive Association of Metabolic Syndrome with a Single Nucleotide Polymorphism of Syndecan-3 (rs2282440) in the Taiwanese Population. Int J Endocrinol 2018, 9282598.

Cornelison, D.D., Filla, M.S., Stanley, H.M., Rapraeger, A.C., and Olwin, B.B. (2001). Syndecan-3 and syndecan-4 specifically mark skeletal muscle satellite cells and are implicated in satellite cell maintenance and muscle regeneration. Dev Biol 239, 79–94.

Cornelison, D.D., Wilcox-Adelman, S.A., Goetinck, P.F., Rauvala, H., Rapraeger, A.C., and Olwin, B.B. (2004). Essential and separable roles for Syndecan-3 and Syndecan-4 in skeletal muscle development and regeneration. Genes Dev 18, 2231–2236.

Cossu, G., Kelly, R., Di Donna, S., Vivarelli, E., and Buckingham, M. (1995). Myoblast differentiation during mammalian somitogenesis is dependent upon a community effect. Proc Natl Acad Sci U S A 92, 2254–2258.

D′Souza, D.M., Al-Sajee, D., and Hawke, T.J. (2013). Diabetic myopathy: impact of diabetes mellitus on skeletal muscle progenitor cells. Front Physiol 4, 379.

De Micheli, A.J., Laurilliard, E.J., Heinke, C.L., Ravichandran, H., Fraczek, P., Soueid-Baumgarten, S., De Vlaminck, I., Elemento, O., and Cosgrove, B.D. (2020). Single-Cell Analysis of the Muscle Stem Cell Hierarchy Identifies Heterotypic Communication Signals Involved in Skeletal Muscle Regeneration. Cell Rep 30, 3583–3595 e3585.

Dumont, N.A., Wang, Y.X., and Rudnicki, M.A. (2015). Intrinsic and extrinsic mechanisms regulating satellite cell function. Development 142, 1572–1581.

El Annabi, S., Gautier, N., and Baron, V. (2001). Focal adhesion kinase and Src mediate integrin regulation of insulin receptor phosphorylation. FEBS Lett 507, 247–252.

Flamini, V., Ghadiali, R.S., Antczak, P., Rothwell, A., Turnbull, J.E., and Pisconti, A. (2018). The Satellite Cell Niche Regulates the Balance between Myoblast Differentiation and Self-Renewal via p53. Stem Cell Reports 10, 970–983.

Fuentealba, L., Carey, D.J., and Brandan, E. (1999). Antisense inhibition of syndecan-3 expression during skeletal muscle differentiation accelerates myogenesis through a basic fibroblast growth factor-dependent mechanism. J Biol Chem 274, 37876–37884.

Ghadiali, R.S., Guimond, S.E., Turnbull, J.E., and Pisconti, A. (2016). Dynamic changes in heparan sulfate during muscle differentiation and ageing regulate myoblast cell fate and FGF2 signalling. Matrix Biol.

Goel, A.J., Rieder, M.K., Arnold, H.H., Radice, G.L., and Krauss, R.S. (2017). Niche Cadherins Control the Quiescence-to-Activation Transition in Muscle Stem Cells. Cell Rep 21, 2236–2250.

Halevy, O., Novitch, B.G., Spicer, D.B., Skapek, S.X., Rhee, J., Hannon, G.J., Beach, D., and Lassar, A.B. (1995). Correlation of terminal cell cycle arrest of skeletal muscle with induction of p21 by MyoD. Science 267, 1018–1021.

Harrington, L.S., Findlay, G.M., and Lamb, R.F. (2005). Restraining PI3K: mTOR signalling goes back to the membrane. Trends Biochem Sci 30, 35–42.

Haruta, T., Uno, T., Kawahara, J., Takano, A., Egawa, K., Sharma, P.M., Olefsky, J.M., and Kobayashi, M. (2000). A rapamycin-sensitive pathway down-regulates insulin signaling via phosphorylation and proteasomal degradation of insulin receptor substrate-1. Mol Endocrinol 14, 783–794.

Huang da, W., Sherman, B.T., and Lempicki, R.A. (2009a). Bioinformatics enrichment tools: paths toward the comprehensive functional analysis of large gene lists. Nucleic acids research 37, 1–13.

Huang da, W., Sherman, B.T., and Lempicki, R.A. (2009b). Systematic and integrative analysis of large gene lists using DAVID bioinformatics resources. Nat Protoc 4, 44–57.

Hwang, J.B., and Frost, S.C. (1999). Effect of alternative glycosylation on insulin receptor processing. J Biol Chem 274, 22813–22820.

Jones, F.K., Hardman, G.E., Ferries, S., Eyers, C.E., and Pisconti, A. (2019). Myoblast Phosphoproteomics as a Tool to Investigate Global Signaling Events During Myogenesis. Methods Mol Biol 1889, 301–317.

Jones, F.K., Stefan, A., Kay, A.G., Hyland, M., Morgan, R., Forsyth, N.R., Pisconti, A., and Kehoe, O. (2020). Syndecan-3 regulates MSC adhesion, ERK and AKT signalling in vitro and its deletion enhances MSC efficacy in a model of inflammatory arthritis in vivo. Sci Rep 10, 20487.

Jones, N.C., Fedorov, Y.V., Rosenthal, R.S., and Olwin, B.B. (2001). ERK1/2 is required for myoblast proliferation but is dispensable for muscle gene expression and cell fusion. J Cell Physiol 186, 104–115.

Klockenbusch, C., and Kast, J. (2010). Optimization of formaldehyde cross-linking for protein interaction analysis of non-tagged integrin beta1. J Biomed Biotechnol 2010, 927585.

Knight, J.D., and Kothary, R. (2011). The myogenic kinome: protein kinases critical to mammalian skeletal myogenesis. Skelet Muscle 1, 29.

Kua, K.L., Hu, S., Wang, C., Yao, J., Dang, D., Sawatzke, A.B., Segar, J.L., Wang, K., and Norris, A.W. (2019). Fetal hyperglycemia acutely induces persistent insulin resistance in skeletal muscle. J Endocrinol 242, M1–M15.

Liu, X., McFarland, D.C., Nestor, K.E., and Velleman, S.G. (2004). Developmental regulated expression of syndecan-1 and glypican in pectoralis major muscle in turkeys with different growth rates. Dev Growth Differ 46, 37–51.

Machackova, K., Chrudinova, M., Radosavljevic, J., Potalitsyn, P., Krizkova, K., Fabry, M., Selicharova, I., Collinsova, M., Brzozowski, A.M., Zakova, L., et al. (2018). Converting Insulin-like Growth Factors 1 and 2 into High-Affinity Ligands for Insulin Receptor Isoform A by the Introduction of an Evolutionarily Divergent Mutation. Biochemistry 57, 2373–2382.

Mashinchian, O., Pisconti, A., Le Moal, E., and Bentzinger, C.F. (2018). The Muscle Stem Cell Niche in Health and Disease. Curr Top Dev Biol 126, 23–65.

Olguin, H., and Brandan, E. (2001). Expression and localization of proteoglycans during limb myogenic activation. Dev Dyn 221, 106–115.

Olguín, H.C., and Olwin, B.B. (2004). Pax-7 up-regulation inhibits myogenesis and cell cycle progression in satellite cells: a potential mechanism for self-renewal. Dev Biol 275, 375–388.

Olguin, H.C., and Pisconti, A. (2012). Marking the tempo for myogenesis: Pax7 and the regulation of muscle stem cell fate decisions. J Cell Mol Med 16, 1013–1025.

Olguin, H.C., Yang, Z., Tapscott, S.J., and Olwin, B.B. (2007). Reciprocal inhibition between Pax7 and muscle regulatory factors modulates myogenic cell fate determination. The Journal of cell biology 177, 769–779.

Perez-Riverol, Y., Csordas, A., Bai, J., Bernal-Llinares, M., Hewapathirana, S., Kundu, D.J., Inuganti, A., Griss, J., Mayer, G., Eisenacher, M., et al. (2019). The PRIDE database and related tools and resources in 2019: improving support for quantification data. Nucleic Acids Res 47, D442–D450.

Peterson, J.M., Bryner, R.W., and Alway, S.E. (2008). Satellite cell proliferation is reduced in muscles of obese Zucker rats but restored with loading. Am J Physiol Cell Physiol 295, C521–528.

Pisconti, A., Banks, G.B., Babaeijandaghi, F., Betta, N.D., Rossi, F.M., Chamberlain, J.S., and Olwin, B.B. (2016). Loss of niche-satellite cell interactions in syndecan-3 null mice alters muscle progenitor cell homeostasis improving muscle regeneration. Skelet Muscle 6, 34.

Pisconti, A., Bernet, J.D., and Olwin, B.B. (2012). Syndecans in skeletal muscle development, regeneration and homeostasis. Muscles, ligaments and tendons journal 2, 1–9.

Pisconti, A., Cornelison, D.D., Olguin, H.C., Antwine, T.L., and Olwin, B.B. (2010). Syndecan-3 and Notch cooperate in regulating adult myogenesis. J Cell Biol 190, 427–441.

Rees, H., Williamson, D., Papanastasiou, A., Jina, N., Nabarro, S., Shipley, J., and Anderson, J. (2006). The MET receptor tyrosine kinase contributes to invasive tumour growth in rhabdomyosarcomas. Growth Factors 24, 197–208.

Reizes, O., Lincecum, J., Wang, Z., Goldberger, O., Huang, L., Kaksonen, M., Ahima, R., Hinkes, M.T., Barsh, G.S., Rauvala, H., et al. (2001). Transgenic expression of syndecan-1 uncovers a physiological control of feeding behavior by syndecan-3. Cell 106, 105–116.

Ronning, S.B., Carlson, C.R., Aronsen, J.M., Pisconti, A., Host, V., Lunde, M., Liland, K.H., Sjaastad, I., Kolset, S.O., Christensen, G., et al. (2020). Syndecan-4(-/-) Mice Have Smaller Muscle Fibers, Increased Akt/mTOR/S6K1 and Notch/HES-1 Pathways, and Alterations in Extracellular Matrix Components. Front Cell Dev Biol 8, 730.

Ruijtenberg, S., and van den Heuvel, S. (2016). Coordinating cell proliferation and differentiation: Antagonism between cell cycle regulators and cell type-specific gene expression. Cell Cycle 15, 196–212.

Shah, O.J., and Hunter, T. (2006). Turnover of the active fraction of IRS1 involves raptor-mTOR- and S6K1-dependent serine phosphorylation in cell culture models of tuberous sclerosis. Mol Cell Biol 26, 6425–6434.

Shin, J., McFarland, D.C., and Velleman, S.G. (2013). Migration of turkey muscle satellite cells is enhanced by the syndecan-4 cytoplasmic domain through the activation of RhoA. Mol Cell Biochem 375, 115–130.

Strader, A.D., Reizes, O., Woods, S.C., Benoit, S.C., and Seeley, R.J. (2004). Mice lacking the syndecan-3 gene are resistant to diet-induced obesity. J Clin Invest 114, 1354–1360.

Tajiri, Y., Kato, T., Nakayama, H., and Yamada, K. (2010). Reduction of skeletal muscle, especially in lower limbs, in Japanese type 2 diabetic patients with insulin resistance and cardiovascular risk factors. Metab Syndr Relat Disord 8, 137–142.

Taylor, J.G.t., Cheuk, A.T., Tsang, P.S., Chung, J.Y., Song, Y.K., Desai, K., Yu, Y., Chen, Q.R., Shah, K., Youngblood, V., et al. (2009). Identification of FGFR4-activating mutations in human rhabdomyosarcomas that promote metastasis in xenotransplanted models. J Clin Invest 119, 3395–3407.

Velleman, S.G., and Song, Y. (2017). Development and Growth of the Avian Pectoralis Major (Breast) Muscle: Function of Syndecan-4 and Glypican-1 in Adult Myoblast Proliferation and Differentiation. Front Physiol 8, 577.

Vuori, K., and Ruoslahti, E. (1994). Association of insulin receptor substrate-1 with integrins. Science 266, 1576–1578.

Whiteford, J.R., Behrends, V., Kirby, H., Kusche-Gullberg, M., Muramatsu, T., and Couchman, J.R. (2007). Syndecans promote integrin-mediated adhesion of mesenchymal cells in two distinct pathways. Exp Cell Res 313, 3902–3913.

Zhang, P., Wong, C., Liu, D., Finegold, M., Harper, J.W., and Elledge, S.J. (1999). p21(CIP1) and p57(KIP2) control muscle differentiation at the myogenin step. Genes Dev 13, 213–224.

