## Supplementary material for "SDC3 acts as a timekeeper of myogenic differentiation by regulating the insulin/AKT/mTOR axis in muscle stem cell progeny": Table S5

**Supplemental Table 5. List of reagents and resources used.**

| REAGENT or RESOURCE | SOURCE | IDENTIFIER |
|---|---|---|
| Antibodies | | |
| Goat anti-SDC3 (For western blotting) | R&D systems | Cat# AF2734; RRID:AB_2183013 |
| Rabbit anti-SDC3 (Immunoprecipitation) | Donated by Prof Alan Rapraeger | N/A |
| Mouse anti-GAPDH | Millipore Sigma | Cat# G8795; RRID:AB_1078991 |
| Rabbit anti-pTyrosine | Cell signaling technology | Cat# 8954; RRID:AB_2687925 |
| Rabbit anti-RPS6 | Cell signaling technology | Cat# 2217; RRID:AB_331355 |
| Rabbit anti-pRPS6$^{S236/S236}$ | Cell signaling technology | Cat# 4858; RRID:AB_916156 |
| Rabbit anti-pRPS6$^{S240/S244}$ | Cell signaling technology | Cat# 5364; RRID:AB_10694233 |
| Rabbit anti-IRS1 | Cell signaling technology | Cat# 3407; RRID:AB_2127860 |
| Rabbit anti-pIRS1$^{S307}$ | Cell signaling technology | Cat# 2381; RRID:AB_330342 |
| Rabbit anti-pIRS1$^{S302}$ | Cell signaling technology | Cat# 2384; RRID:AB_330360 |
| Rabbit anti-AKT | Cell signaling technology | Cat# 9272; RRID:AB_329827 |
| Rabbit anti-pAkt$^{S473}$ | Cell signaling technology | Cat# 4060; RRID:AB_2315049 |
| Rabbit anti-ERK1/2 | Cell signaling technology | Cat# 9102; RRID:AB_330744 |
| Rabbit anti-pERK1/2$^{T202/Y204}$ | Cell signaling technology | Cat# 9101; RRID:AB_331646 |
| Rabbit anti-Insulin receptor β | Cell signaling technology | Cat# 3025; RRID:AB_2280448 |
| Rabbit anti-IGF1 receptor β | Cell signaling technology | Cat# 9750; RRID:AB_10950969 |
| Mouse anti-Myosin heavy chain | DSHB | Cat# MF 20; RRID:AB_2147781 |
| Normal-rabbit IgG | Cell signaling technology | Cat# 2729; RRID:AB_1031062 |
| Chemicals and Recombinant Proteins | | |
| Capivasertib (AKT inhibitor) | Selleckchem | Cat# S8019 |
| Insulin | Sigma | Cat# I0516 |
| FGF2 | Donated by Prof David Fernig | N/A |
| DAPI | ThermoFisher Scientific | Cat# D1306 |
| Lipofectamine 2000 transfection reagent | ThermoFisher Scientific | Cat# 11668019 |
| Heparinase III | IBEX | Cat# 50-012-001 |
| Collagenase type I | Worthington Biochemical | Cat# LS004177 |
| Puromycin | Millipore Sigma | Cat# P8833 |
| Chondroitinase ABC | Millipore Sigma | Cat# C3667 |
| Commercial kits | | |

| | | |
|---|---|---|
| Pierce BCA Protein Assay Kit | ThermoFisher Scientific | Cat# 23227 |
| cOmplete™ Protease Inhibitor Cocktail | Millipore Sigma | Cat# 4693116001 |
| PhosSTOP | Millipore Sigma | Cat# 4906845001 |
| ECL™ Prime Western Blotting Detection Reagent | Millipore Sigma | Cat# GERPN2236 |
| Experimental Models: Cell Lines | | |
| Mouse: C2C12 cells | ATCC | ATCC Cat# CRL-1772; RRID:CVCL_0188 |
| Mouse: Ctrl cells | This paper | N/A |
| Mouse: $S3^{kd}$ cells | This paper | N/A |
| Experimental Models: Organisms/Strains | | |
| Mouse: wild type: C57Bl/6J | Charles River Laboratory, UK | C57Bl/6J |
| Mouse: *Sdc3*$^{-/-}$ | Reizes et al. (2001) | N/A |
| Oligonucleotides | | |
| siRNA targeting sequence: INSR #1: SASI_Mm01_00090856 | Millipore Sigma | Cat# NM_010568 |
| siRNA targeting sequence: INSR #2: SASI_Mm01_00090859 | Millipore Sigma | Cat# NM_010568 |
| siRNA scrambled control #1 | Millipore Sigma | Cat# SIC001 |
| Recombinant DNA | | |
| Control knockdown plasmid: pLKO mouse | Millipore Sigma | Cat# SHC001 |
| Sdc3 knockdown plasmid: pLKO mouse shRNA SDC3, TRCN0000071990 | Millipore Sigma | Cat# SHCLND-NM_011520 |
| Software and Algorithms | | |
| Prism 8 | GraphPad Prism | RRID:SCR_002798 |
| Ingenuity Pathway Analysis | Ingenuity Pathway Analysis | RRID:SCR_008653 |
| DAVID, functional annotation tool | Huang da et al. (2009) | DAVID; RRID:SCR_001881 |
| Progenesis QI | Nonlinear Dynamics | http://www.nonlinear.com/progenesis/qi/ |
| PEAKS studio 8; PTM | Han et al. (2011) | https://www.bioinfor.com/ |
| FIJI | Schindelin, J *et al.* (2012) | RRID:SCR_002285 |
| Bayesian Markov chain Monte Carlo simulation | This paper | https://github.com/PGB-LIV/JonesSDC3PhosphoproteomicsPaper |
| FIJI script used for quantification of nuclei and myofiber size | Arecco *et al.* (2016) | https://github.com/Piscontilab/Fiji-script |


Han, X., He, L., Xin, L., Shan, B., and Ma, B. (2011). PeaksPTM: Mass spectrometry-based identification of peptides with unspecified modifications. J Proteome Res *10*, 2930-2936.

Huang da, W., Sherman, B.T., and Lempicki, R.A. (2009). Systematic and integrative analysis of large gene lists using DAVID bioinformatics resources. Nat Protoc *4*, 44-57.

Reizes, O., Lincecum, J., Wang, Z., Goldberger, O., Huang, L., Kaksonen, M., Ahima, R., Hinkes, M.T., Barsh, G.S., Rauvala, H., *et al.* (2001). Transgenic expression of syndecan-1 uncovers a physiological control of feeding behavior by syndecan-3. Cell *106*, 105-116.
