## Supplemental Figures for "SDC3 acts as a timekeeper of myogenic differentiation by regulating the insulin/AKT/mTOR axis in muscle stem cell progeny"

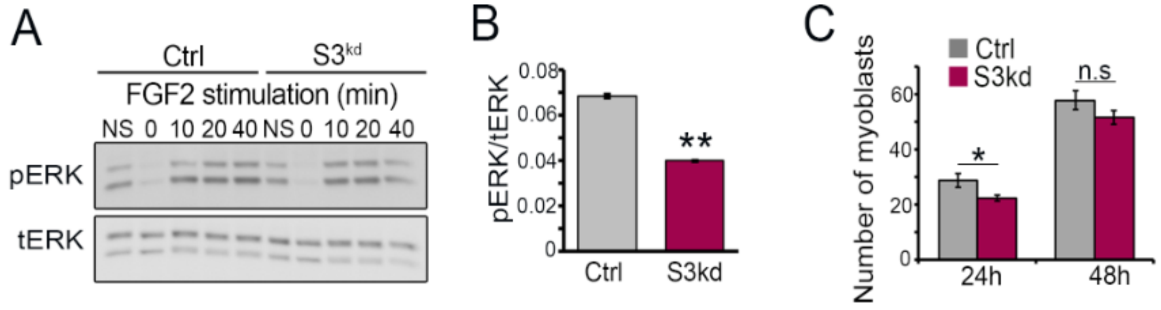


**Figure S1. SDC3-depleted cells recapitulated phenotypes observed in *Sdc3*$^{-/-}$ myoblasts.**

(A) Control (Ctrl) and SDC3-depleted (S3$^{kd}$) cells were cultured in in growth medium, then serum-starved for 5 hours before stimulating with 2nM FGF2 for the indicated time points. Cells were lysed and lysates subjected to western blotting. Levels of phosphorylated ERK (pERK) and total-ERK (tERK) were measured. NS, non-starved. N=3

(B) Quantification of serum-starved Ctrl and S3$^{kd}$ cells from (A). N=3

(C) Equal numbers of Ctrl and S3$^{kd}$ cells were cultured in growth medium for 24 and 48 hours. Cells were stained with DAPI to visualise nuclei and and the number of myoblasts per image taken was counted using a bespoke FIJI script. Error bars represent standard error. *$p<0.05$. n.s, non-significant. Averages from three independent experiments were plotted, where 18-20 images were taken and analyzed per replicate (total N = minimum 54 images per data point).

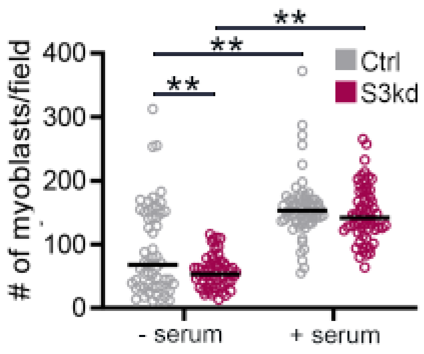


**Figure S2. SDC3-depleted cells have a reduced capacity to adhere to laminin.**

Equal numbers of control (Ctrl) and SDC3-depleted (S3kd) cells were seeded onto laminin-coated wells and cultured for 1 hour under normal growth conditions. Cells were washed gently once with PBS to remove non-attached cells. Cells were then fixed with PFA and stained with DAPI to visualise nuclei. The number of cells per image were counted using a bespoke FIJI script. Averages from three independent experiments were plotted, where 16-20 images per replicate were taken and analyzed (total N = minimum 48). **$p<0.01$.
